# Divergence and convergence in epiphytic and endophytic phyllosphere bacterial communities of rice landraces

**DOI:** 10.1101/2024.06.29.601313

**Authors:** Pratibha Sanjenbam, Deepa Agashe

## Abstract

Phyllosphere-associated microbes can significantly alter host plant fitness, with distinct functions provided by bacteria inhabiting the epiphytic (external surface) vs. endophytic niches (internal leaf tissue). Hence, it is important to understand the assembly and stability of these phyllosphere communities, especially in field conditions. Broadly, epiphytic communities should encounter more environmental changes and immigration, whereas endophytic microbiomes should face stronger host selection. We analysed the structure and stability of leaf phyllosphere microbiomes of four traditionally cultivated rice landraces and one commercial variety from northeast India grown in the field for three consecutive years, supplemented with opportunistic sampling of 8 other landraces. Epiphytic and endophytic bacterial communities shared dominant core genera such as *Methylobacterium* and *Sphingomonas*. Consistent with an overall strong environmental effect, both communities varied more across sampling years than across host landraces. Seeds sampled from a focal landrace did not support vertical transmission of phyllosphere bacteria, suggesting that both types of communities are assembled anew each generation. Despite these points of convergence, epiphytic communities had distinct composition and significantly higher microbial load, and were more rich, diverse, modular, and unstable than endophytic communities. Finally, focused sampling of one landrace across developmental stages showed that the divergence between the two types of communities arose primarily at the flowering stage. Thus, our results show both convergent and divergent patterns of community assembly and composition in distinct phyllosphere niches in rice, identifying key bacterial genera and host developmental stages that may aid agricultural interventions to increase rice yield.

## INTRODUCTION

The phyllosphere is the aerial part of a plant and is home to various microbes in distinct niches. For instance, the leaf phyllosphere has microbes residing either on the surface (epiphytes) or inside the leaf tissue (endophytes). These leaf-associated microbes (henceforth “phyllosphere microbes”) can greatly impact host plant fitness, ranging from nutrient acquisition to resistance against pathogens (reviewed in (1–4). Hence, understanding the assembly and maintenance of phyllosphere microbiomes is an important research focus. Broadly, the phyllosphere microbiome is shaped by a combination of processes: ecological drift, host selection, environmental factors, dispersal, and evolution (5). Though patterns of host genotype- or population-specific phyllosphere microbiomes ((3, 6–10) are consistent with a stronger host selection effect, in many plants the phyllosphere microbiome is influenced by both host plant genotype and abiotic environmental factors such as light, temperature, humidity, and water (11–13). Recent work also suggests that effects of host vs. environment can be dynamic. In an evolution experiment with tomato plants inoculated with naturally derived phyllosphere communities, both host genotype and environment had strong effects on microbial community structure, but the host genotype effect weakened across passages (14). Typically, we expect strong host selection to reduce stochasticity in community assembly, whereas strong environmental effects are expected to introduce more variation across microbiomes of conspecific plants. Indeed, temporal (including seasonal) variation in phyllosphere microbiome is commonly observed across multiple plant species (13, 15–17). However, most prior studies have focused on the epiphytic community, with far fewer analyses of the impact of host vs. environmental factors on the leaf endosphere.

The epiphytic and endophytic plant niches are quite distinct and impose different selection pressures on colonizing microbes (1). Bacteria in each niche thus exhibit different suites of traits, provide distinct benefits to the host, contribute differentially to global nutrient cycles, and respond in distinct ways to climate change (9, 10, 18). An interesting example comes from the pathogens *Pseudomonas syringiae* which transitions from an early epiphytic commensal phase to a later pathogenic endophytic stage, with associated changes in suites of key genes expressed in each stage (19). Colonization of epiphytic vs. endophytic niches also involves different host vs. environment effects and ecological processes governing community assembly. For instance, we expect that the endophytic community should experience stronger host selection, with a relatively weaker influence of external environmental factors. In contrast, we expect the epiphytic community to be more strongly affected by the environment because it is continually and directly exposed to a large influx of microbes from several sources such as soil, air, and water. Additionally, local dispersal of microbes from neighboring plants can contribute to the assembly of the phyllosphere epiphytic microbial community (20) (Meyer et al 2022). Lastly, we also expect to see distinct communities and core membership across different plant niches (18)(Trivedi et al 2020). However, very few studies have directly tested for divergence in community assembly and maintenance of epiphytes vs. endophytes. The few studies that have been conducted report variable results. In the endophytic microbial community associated with Acacia (13)and rice plants (21), a stronger host effect was observed than for epiphytes; host effects were similar for endophytic and epiphytic communities of olive trees (Mina et al 2020); whereas in sugar maples, the host effect was greater in epiphytic than the endophytic phyllosphere community (22). Thus, the effect of host vs. environment may not be easily generalizable, and it is unclear whether the variability arises due to differences in host species, or differences and limitations in sampling and experimental design (e.g., sampling only one plant growth stage). Hence, a deeper understanding of differences between epiphytic and endophytic phyllosphere microbiomes requires monitoring of the dynamics of both phyllosphere microbiomes across the same host species, and across time and plant growth stages.

Another important reason to compare epiphytic vs. endophytic communities is to better understand variation in core community members that may be most relevant for plant fitness. A large body of work has demonstrated important functions provided by core phyllosphere bacteria, including nitrogen fixation, resistance against pathogens, and drought tolerance (4, 23). Bacteria from the genera *Methylobacterium* and *Sphingomonas* from alpha-proteobacteria are well-known examples of beneficial symbionts for many plants (24–26), enhancing seed germination (27, 28), promoting plant growth (26, 29, 30) and providing resistance against pathogens (31, 32) Additionally, we recently demonstrated substantial yield benefits from a specific *Methylobacterium* strain on its host rice plants in field conditions, though the effect of *Methylobacterium* inoculation was not generalizable across host-bacterium pairs (33). The impacts of specific bacteria may differ not only across hosts, but also in a community context, and with environmental variation. For instance, inoculation of the phyllosphere microbiome on tomato plants confers resistance against the pathogen *Pseudomonas syringae*, but this effect is contingent on available nutrient resources (34). More generally, the action of specific microbes can either facilitate or hinder the success of subsequent colonizers (35). However, in such studies it is difficult to identify which community members provide the observed beneficial function. One way to begin approaching this problem is to identify sets of core microbes that consistently co-occur at high abundance, despite temporal variation at the community level. Such co-occurring sets of taxa may indicate interacting partners that respond to host-imposed selection, and testing for unique co-occurring taxa across plant niches may thus allow the identification of niche-specific, putatively interacting bacteria.

We addressed these gaps using multiple rice landraces grown in the state of Manipur in north-east India. These local rice varieties have been selected for specific traits (such as yield, aroma, and grain color) for many generations, resulting in substantial phenotypic and genotypic diversity (36–39). We surmised that this host diversity might drive differential selection on bacterial taxa associated with each landrace, and we quantified phyllosphere community dynamics in leaf niches across time. We collected leaf samples from four rice landraces and one commercial high-yielding variety (HYV) across three consecutive growing seasons (Fig S1A). Using 16S rRNA amplicon sequencing, we jointly analysed leaf epiphytic and endophytic communities. Our work thus quantifies the impacts of various factors in shaping phyllosphere communities, and provides a deeper understanding of host-plant-associated bacterial communities in the field.

## METHODS

### Sample collection

We collected leaf samples of flowering rice plants from multiple sites in Manipur in 2019, 2020, and 2021 (Table S1). We focused on 4 phenotypically distinct rice landraces that were consistently grown by the farmer during these years, i.e., Chakhao (CK), Phouren-mubi (PM), Phoungang (PN), and Tolenphou (TP), and one commercially developed high yielding rice variety (HYV) (Table S1). For each landrace, we randomly chose 5–10 plants from the field, cut a 3–4 cm length of the flag leaf from each plant, and immediately immersed it in 1mL of 1X Redford buffer (1M tris HCl, 0.5M EDTA, 1.24% Triton) placed on ice, as described earlier (33)(Sanjenbam et al 2022) (Fig S1A). In 2021, we similarly collected samples of leaves from different plant growth stages from landrace CK (Fig S1B). Finally, in 2019 and 2020, we sampled seeds from landrace CK. After harvesting, seeds were transported in plastic bags to the laboratory, where they were stored at –80°C.

### Determining phyllosphere and seed microbiome composition

We followed our previously described protocol to extract DNA for epiphytic bacterial communities (33). Briefly, leaf samples were processed immediately after transport to the laboratory for epiphytic DNA extraction. Tubes were placed horizontally on a vortex for 2 hrs with intermittent shaking to dislodge surface bacteria, after which we removed the leaves. We centrifuged tubes at 10,000 rpm for 25 mins and used the pellet for extracting DNA as per the Qiagen DNeasy blood and tissue kit instructions. To extract DNA from endophytic communities, we used the same leaf sample used for epiphytes, washed it once with PBS, and crushed it using liquid nitrogen. We used the powdered leaf (∼100mg) for DNA extraction using the same kit as above, with the following modifications. We added 180µL of lysis buffer and 20µL proteinase K and incubated at 56°C overnight. To remove leaf debris, we used a brief centrifugation step and used the supernatant for subsequent steps as recommended in the kit instructions. We used 30 µL of nuclease-free water for eluting DNA. We note that 2019 samples were processed using a different kit (PowerSoil DNA isolation kit, Mo Bio). For seed microbiomes, we extracted epiphytic DNA using 3 seeds/replicate/landrace as described above, in 500µL Redford buffer. For seed endophytic bacteria, we removed the seed husk and surface-sterilized seeds using immersion in 70% ethanol. Sterilized seeds were crushed using liquid nitrogen and 100mg of tissue was used for DNA extraction.

We quantified the DNA concentration for all samples using Qubit, and then amplified the V3-V4 hypervariable region of the 16S rRNA using standard Illumina primers (40). We used PNAs (polypeptide nucleic acid) to block the amplification of plant mitochondrial and chloroplast DNA in our 16S rRNA PCR, as described previously (41). Finally, we carried out amplicon sequencing on the Illumina Miseq platform (300×2 paired-end).

### qPCR to quantify the total bacterial load

We used qPCR to quantify the total bacterial load for 2021 samples of the four focal landraces (CK, PM, PN, and TP) for both epiphytes and endophytes, and across different growth stages of landrace CK in the same year. We designed 16S rRNA primers (forward: 5’-CGGTAATACGGAGGGTGCAA-3’; reverse: 5’-TCCTCCAGATCTCTACGCAT -3’) to selectively amplify bacterial DNA (200bp). We used primers for the rice gene Os*ACTIN1* (forward:5’-ATGAAGTGCGACGTGGATATTAG-3’; reverse:5’GGGCGACCACCTTGATCTTC-3’) to estimate the amount of host DNA for both epiphytic and endophytic samples. We used 1µL of DNA as a template for qPCR (BioRad CFX system), testing 5 biological replicates/landrace/community type. For each biological replicate, we carried two technical replicate qPCR reactions and normalized Ct values for the target gene against *OsACTIN1* to obtain the normalized total bacterial load as 2^-ΔCt^, where ΔC_t_ = C_t(16S rRNA gene)_ – C_t(actin)_. This quantity approximates the amount of bacterial DNA per unit host DNA.

### Statistical analysis

We analyzed and visualized data using R (42). To analyse amplicon sequencing data, we used the DADA2 workflow (43). After filtering out chloroplast and mitochondrial reads, we obtained an average of 50,000 reads (range 10,000 to 500,000) per epiphytic sample and an average of 29,000 reads (range 5,000 to 68,000) per endophytic sample (Fig S2A–B). Our negative controls without template had an average of 98 reads, indicating low levels of reagent contamination. Rarefaction analysis of a subset of samples showed that ∼20,000 reads/sample for epiphytic samples and ∼10,000 reads/sample for endophytic samples leads to near-saturation of community richness (Fig S2C). Several endophyte samples initially had low read depth, and this problem was not always solved by additional sequencing. Samples that had fewer reads than the threshold noted above were excluded from further analysis. Bacterial taxa with <1% of the total reads were combined into a single bin as “rare taxa” or “others”, only for visualization. For all statistical analyses, the full matrix of relative abundances of each taxon was used, unless specified otherwise.

To quantify the impact of different factors (e.g., landrace and sampling year) on community composition, we first calculated Bray-Curtis distances between the samples, and carried out PERMANOVA (Permutational analysis of variance) using the function ‘adonis2’ in the package ‘vegan’ (44). We used this distance matrix to carry out Canonical Analysis of Principal Coordinates based on Discriminant Analysis (CAPdiscrim) (45) using the R package ‘Biodiversity R’ (46), permuting 1000 times. Two of the dominant linear discriminants (LD) were plotted to visualize each cluster with ellipses reflecting 95% confidence intervals using the function ‘ordiellipse’ in the R package ‘vegan’ (44). We used the package ‘ggplot2’ (47) for visualization.

To compare the richness and Shannon diversity index for epiphytes vs. endophytes, we carried out paired t-tests. To estimate the effect of landrace in a given year, we compared microbiomes from all the landraces collected in that year. To estimate the effect of sampling year, we compared microbiomes for the 4 focal landraces across all years. We also used ANOVA for testing the interaction effect of landrace, year, community type or growth stage on both epiphytic and endophytic microbial communities. We used the R package NST (48) to measure taxonomic beta diversity (β_rc-_ Raup-Crick Dissimilarity) as a measure of the degree of individual variation across community types in the focal rice landraces (49). This metric ranges from -1 to +1, with values of +1 indicating that communities are more dissimilar across individuals, and values of -1 indicating more similar communities across individual samples (49) (in our case, each sample was from a different individual plant).

For network analysis, we first used the relative abundance microbiome dataset (4 focal landraces for 3 years) to generate a correlation matrix by calculating all pairwise Spearman’s rank correlations for all bacterial genera (combining data for multiple ASVs assigned to a given genus) across all epiphytic as well as endophytic samples. We filtered out bacterial pairs where the absolute correlation coefficient was less than 0.5 and p>0.05, retaining only significant pairs to construct the occurrence network of bacterial taxa. Finally, we used Gephi (50) for visualizing the network.

## RESULTS

Since epiphytic communities are more exposed to the environment, we expect greater stochasticity in epiphytic community assembly and maintenance, but more deterministic processes to govern the endophytic community. Accordingly, we outlined the following expectations regarding differences between epiphytic and endophytic communities:

1. Higher bacterial richness and diversity in epiphytic communities, in all landraces and years.
2. Weaker effect of host rice landrace and stronger effect of sampling year on epiphytic community composition.
3. Lower consistency and stability of epiphytic communities across landraces and years.

### Epiphytic and endophytic phyllosphere communities converge on similar core genera in all host landraces

Before testing the specific hypotheses outlined above, we describe the broad structure and composition of the phyllosphere communities. As expected from prior work (2, 3, 51), the phyllosphere communities of all four traditional rice landraces were dominated by the genera *Methylobacterium, Sphingomonas, Pantoea, Aureimonas*, and *Pseudomonas* (Fig 1). Remarkably, both epiphytic and endophytic communities showed the same dominant taxa, and this was also true for the high-yielding variety (Fig S3A) and for the eight additional rice landraces that were collected sporadically (Fig S3 B-D). The only exception was the landrace Moirangphou (MP), where *Methylobacterium* was not dominant and instead, *Rhodococcus* and *Acinetobacter* had higher abundance (Fig S3B). Of the dominant taxa, *Methylobacterium* was the single most abundant genus (on average ∼20–40% of all reads in epiphytic communities and 10–70% in endophytic communities) (Fig 1, Fig S3). At a lower taxonomic level, 4 *Methylobacterium* ASVs (ASV1, ASV3, ASV6, and ASV12) accounted for ∼10–30% of the total *Methylobacterium* abundance as epiphytes and endophytes, and may be potentially important strains for the rice plants (Fig S4). These patterns were also observed in a core microbiome analysis that accounts for both prevalence (across samples) and abundance of community members (Fig 2). *Methylobacterium and Sphingomonas* were both dominant and consistently present across the four focal landraces (∼98% prevalence in samples with high relative abundance, Fig 2). However, *Pantoea* and *Aureimonas* were relatively less prevalent and typically occurred at lower relative abundance (∼50% prevalence across the samples with low relative abundance), in epiphytic and endophytic communities respectively. Thus, at the level of core community membership, the epiphytic and endophytic communities were remarkably convergent across individual plants, as well as across years and host landraces.

**Fig 1:**
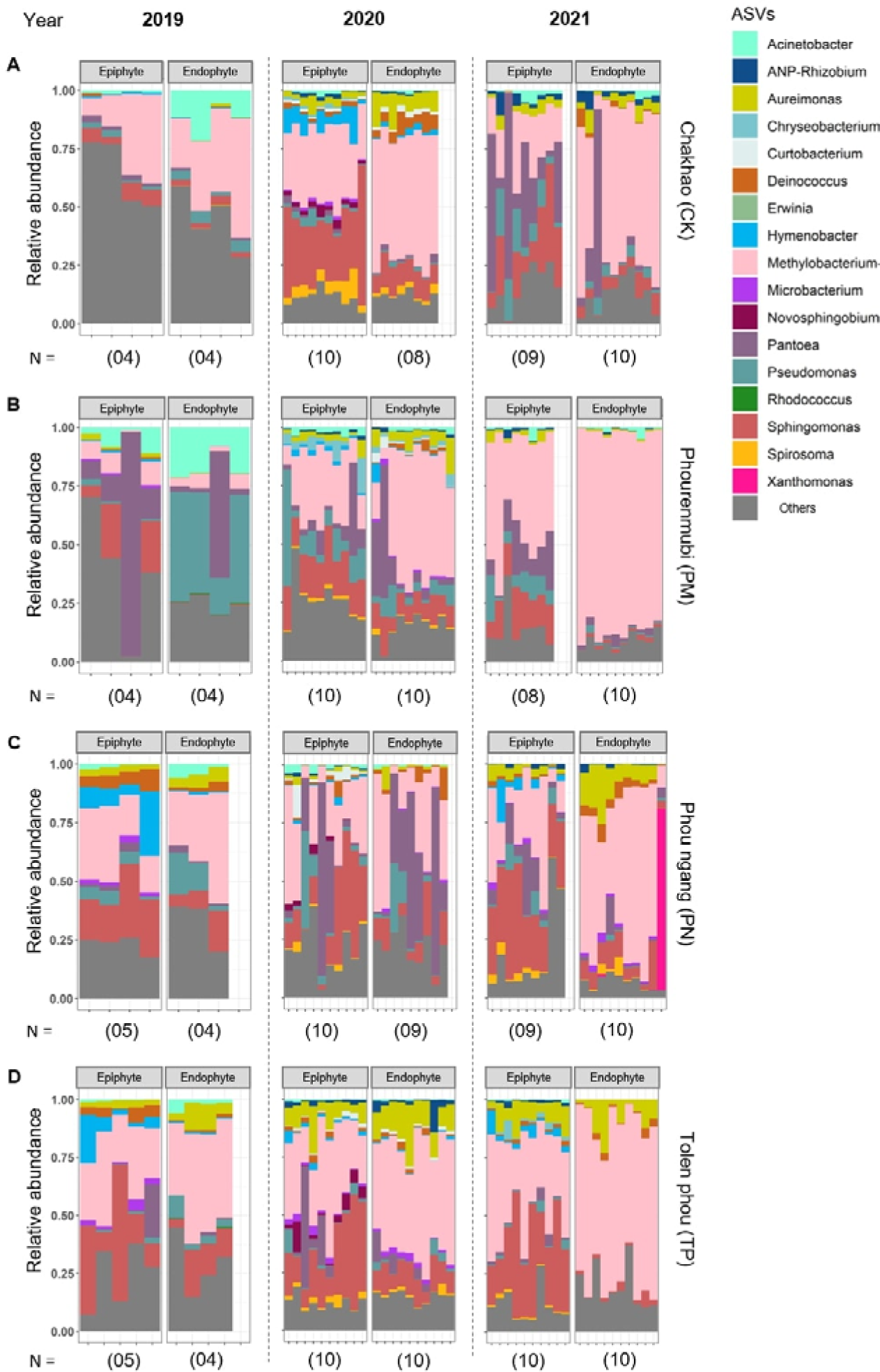
Dominant taxa in phyllosphere microbiomes. Stacked bar plots show the relative abundance of bacterial ASVs (genus level) for each landrace sampled in each year (A) Chakhao (CK); (B) Phouren-mubi (PM); (C) Phou ngang (PN); (D) Tolenphou (TP). Sample size (number of replicate plants) is indicated in parentheses. Taxa with <1% reads, or that were not assigned to known bacteria in the database, were combined into “others”.

**Fig 2:**
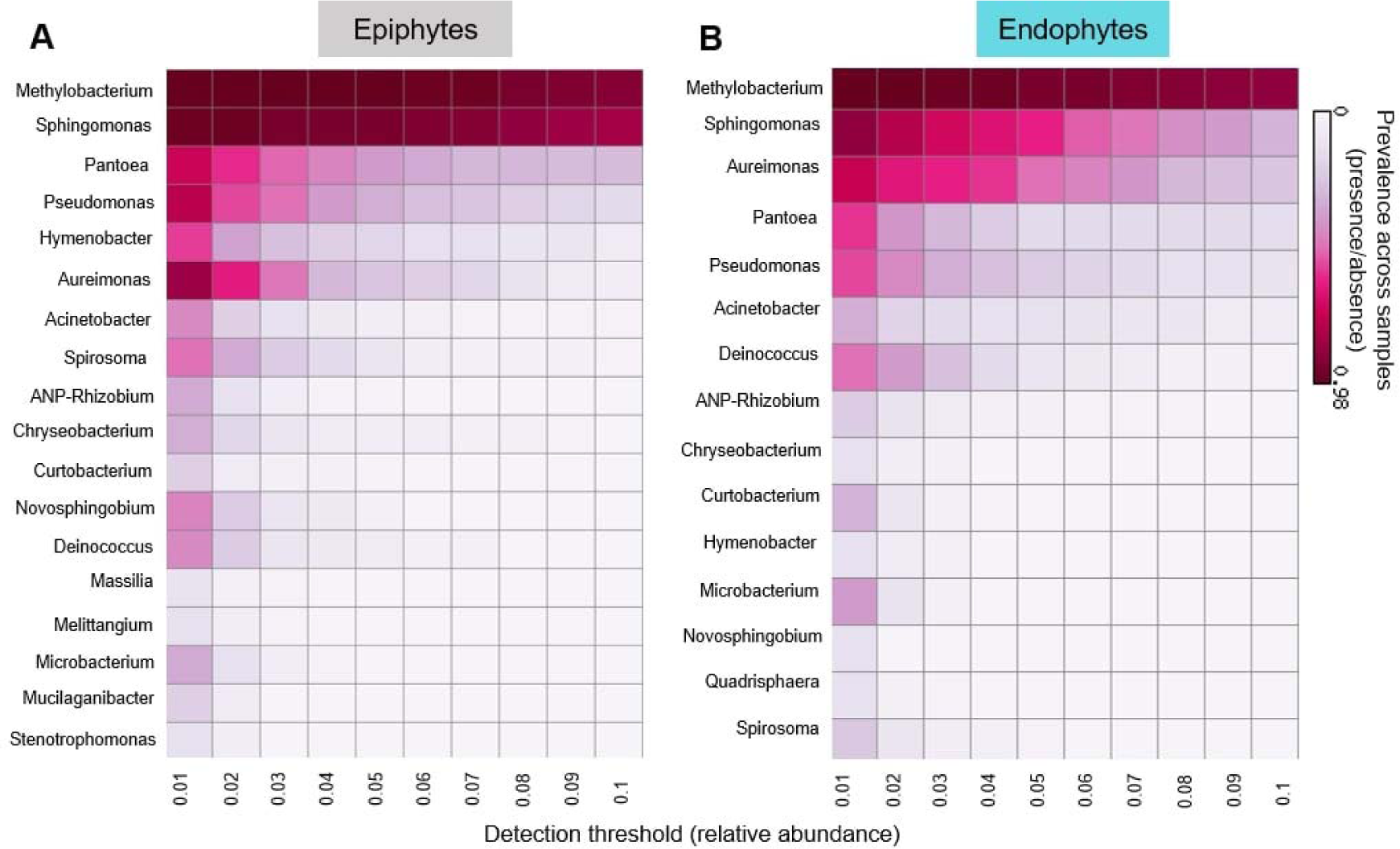
Similar core genera comprise epiphytic and endophytic phyllosphere communities. Heatmaps show the prevalence of core bacterial genera in (A) epiphytic and (B) endophytic communities. Prevalence was calculated as the % of all samples where a given taxon was present with the indicated threshold of relative abundance. For instance, *Methylobacterium* is prevalent (found in most samples) across a wide range of relative abundance (0.01 to 0.1), whereas *Pantoea* is prevalent only when rare.

### Phyllosphere epiphytic bacterial communities tend to be more diverse, rich, and abundant than endophytic bacterial communities

Using the microbiome data and focusing on the four traditionally cultivated rice landraces, we next tested our specific predictions about differences between epiphytic and endophytic communities. Overall, microbiome composition across individual plants from a given landrace in a given year tended to be similar (Fig 1, Fig S3). However, in all cases, epiphytic communities were significantly more dissimilar across individual plants, compared to endophytic communities (Fig 3), supporting our prediction that the epiphytic community is more vulnerable to stochastic processes. Epiphytic bacterial communities of all landraces also tended to be more diverse than endophytic communities in all years (Fig 4A; also see Fig S5), though the difference was not always statistically significant (results of paired t-tests comparing epiphytic vs. endophytic communities for a given landrace and year are shown in Fig 4A and Fig S5; Kruskal Wallis tests for Shannon diversity ∼ community type in each year: 2019, p = 0.004, 2020, p = 1.2 x 10^-5^; 2021, p = 5 x 10^-6^). Similarly, epiphytic communities tended to have higher richness in all cases (Fig 4B, Fig S5). Again, the landrace Moirangphou (MP) was exceptional, showing an opposite pattern in 2020, with higher diversity and richness in endophytic communities (Fig S5). Together, these results support our first prediction that epiphytic communities should have higher richness and diversity.

**Fig 3:**
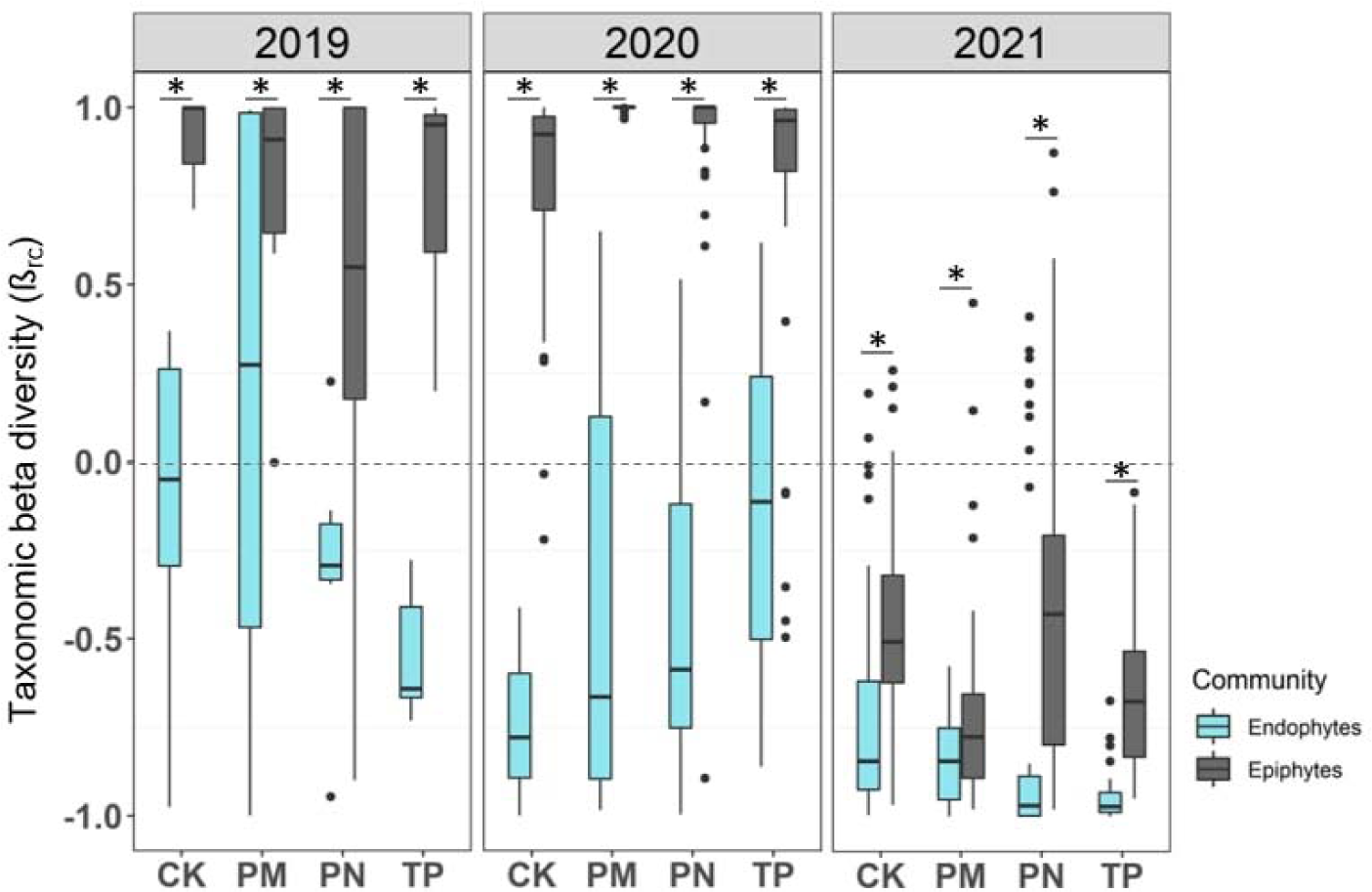
Individual variation is higher in the epiphytic than the endophytic phyllosphere community. Boxplots show taxonomic beta diversity (β_rc_) of bacterial communities of rice landraces across years. Asterisks indicate significant differences across epiphytic and endophytic niches (Wilcoxon test, p<0.05). Sample sizes for the year 2019 range from 4-5 biological replicates/landrace; 2020 range from 7-10 biological replicates/landrace; 2021 range from 8-10 biological replicates/landrace (Table S1).

**Fig 4:**
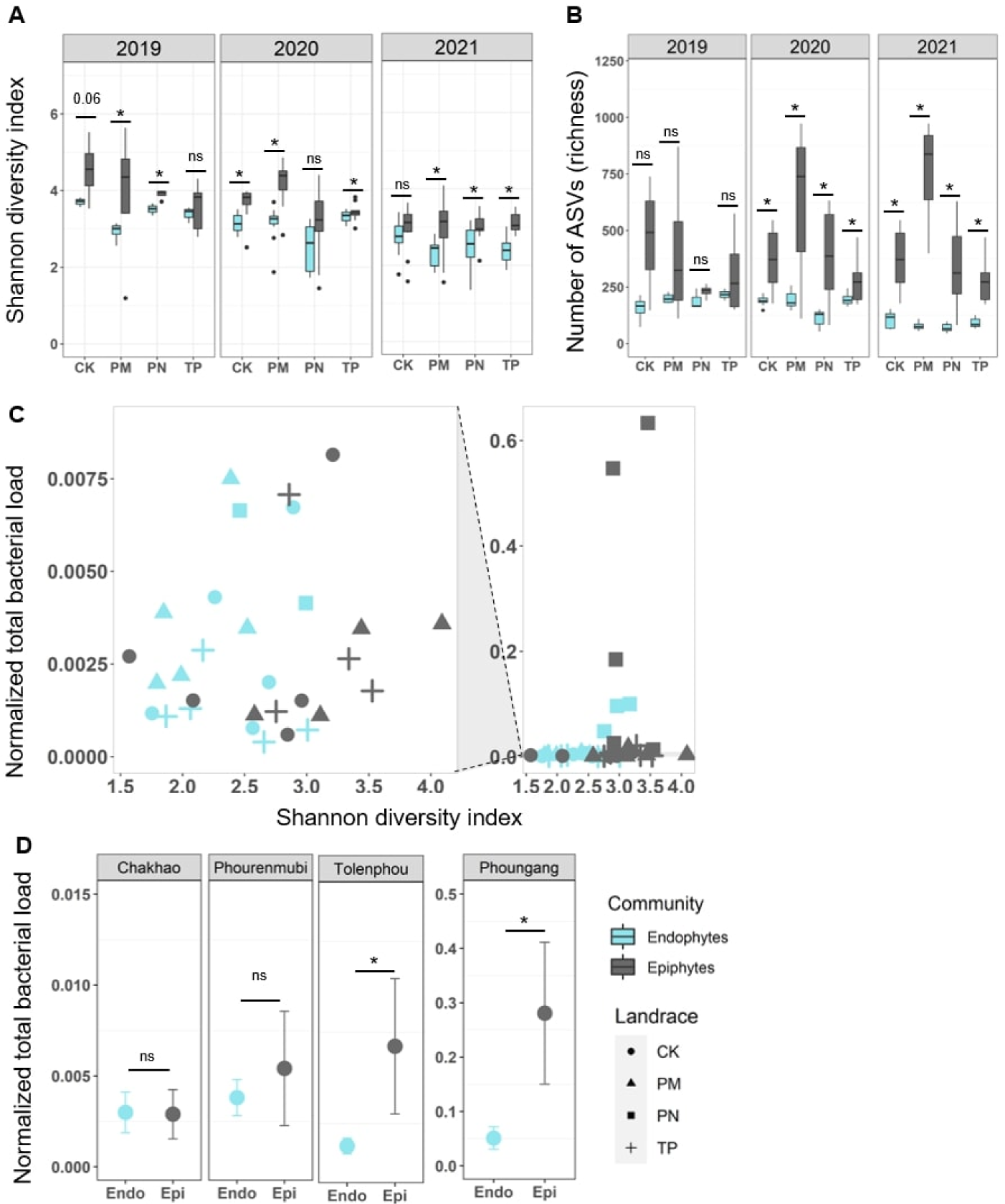
Epiphytic microbial communities are more diverse and abundant than endophytic communities. Boxplots show (A) Shannon’s diversity and (B) the richness of bacterial communities of rice landraces across years. Asterisks indicate p<0.05 (paired t-tests). Sample sizes for the year 2019 range from 4-5 biological replicates/landrace; 2020 range from 7-10 biological replicate/landrace; 2021 range from 8-10 biological replicates/landrace (Table S1). (C) Shannon diversity of communities as a function of the normalized total bacterial load. The right-hand panel shows the full dataset that includes some outliers (squares); the left-hand plot represents a truncated y-axis to highlight variation in the bulk of the dataset, excluding the potential outliers. Each point represents one biological replicate plant (n=5/landrace), for each of the four focal rice landraces sampled in 2021. (D) Normalized total bacterial load (bacterial DNA/host DNA) for phyllosphere communities in a given landrace and year 2021 (asterisks indicate p<0.05, paired Wilcox tests) (ANOVA, load ∼ landrace x community, p _landrace_ = 0.005; p _community_ = 0.048, p _landrace x community_ = 0.03).

Did community diversity and richness reflect total bacterial load, i.e., success of colonization of the phyllosphere? We tested this using samples collected in 2021. We did not find a correlation between bacterial load and either Shannon diversity (epiphytes, Spearman’s rho = 0.33, p = 0.14; endophytes, Spearman’s rho = 0.35, p = 0.12; Fig 4C) or richness (Fig S6), suggesting that community diversity and richness do not necessarily correspond with bacterial colonization success. For instance, despite substantially larger microbial loads in the landraces Tolenphou (TP) and Phoungang (PN), bacterial diversity was similar across landraces in 2021, suggesting that the same taxa were colonizing more successfully in these two landraces. However, bacterial loads did vary substantially across landraces, with Phoungang (PN) showing the highest loads for the epiphytic community — an order of magnitude higher than other landraces (Fig 4D). Additionally, epiphytic communities had higher total loads than endophytes (Fig 4D), potentially reflecting greater exposure to the environment and weaker selection from the host. Thus, patterns of variation in microbiome community composition (diversity and richness), as well as load, broadly support our first prediction of greater stochasticity and environmental impacts on epiphytic communities.

### Sampling year has stronger effects than the host landrace on both epiphytic and endophytic communities

Next, we tested our second prediction, that host rice landrace identity should have a stronger impact on endophytic compared to epiphytic bacterial communities, whereas sampling year should have a stronger effect on epiphytic communities. Overall, both landrace and sampling year influenced the richness and diversity of both sets of communities. We found significant impacts of landrace and marginally significant effects of landrace x year interaction for both epiphyte and endophyte community richness, but a significant effect of sampling year only for endophytes (ANOVA, richness ∼ landrace x year: for epiphytes, p _landrace_ = 2.8×10^-9^, p _year_ = 0.15, p _landrace x year_ = 0.047; for endophytes, p _landrace_ = 2.9×10^-5^, p _year_ < 2×10^-16^, p _landrace x year_ = 0.05). In the case of community diversity, we found significant variation only across years but not across landraces for epiphytes (ANOVA, Shannon diversity index ∼ landrace x year: p _landrace_ = 0.18, p _year_ = 1.3×10^-6^, p _landrace X year_ = 0.2), whereas for endophytes, both landrace and year had significant impacts (ANOVA, Shannon diversity index ∼ landrace x year, p _landrace_ = 0.03, p _year_ = 1.3×10^-10^, p _landrace X year_ = 0.6). Overall, both landrace identity and sampling year influenced endophytes more consistently than epiphytes, suggesting that epiphytic diversity and richness may also vary as a function of other unknown factors.

We then tested the impact of landrace and sampling year on community composition. We visualized the effect of landrace using linear discriminant analysis, which showed clustering of both epiphytic and endophytic microbial communities across years (Fig S7). As expected, sampling year had a higher impact on epiphytic communities compared to endophytes, explaining 20% and 16% of variation respectively (PERMANOVA, Table 1A). Similarly, as expected, host landrace had a higher impact on endophytic communities, but the difference was small (8% in epiphytic communities and 10% in endophytic communities, Table 1A). On including two additional landraces sampled in 2 of the 3 years and the high-yielding variety, the effect of the host landrace on both epiphytic and endophytic communities became stronger (12% and 15% respectively, Table S2). However, this was primarily driven by Moirangphou (MP), which had a very distinct community composition, as described above (Fig S3). Within a given year, the effect of host landrace was generally much higher, ranging from 20–30% for epiphytic communities and 29–45% for endophytic communities (Table S2). Together, these results provide some support for our second prediction, though the magnitude of differences between epiphytic and endophytic communities was low.

**Table 1:** Effect of landrace, year, and community type on phyllosphere microbial communities. Tables show the output of PERMANOVA analysis of communities from (A) the four focal landraces and (B) all landraces. Significant p-values are represented in bold.

| **A** |  | Landrace |  | Year |  | Landrace x Year |  |
| --- | --- | --- | --- | --- | --- | --- | --- |
| Community type | Year | R<sup>2</sup> | p | R<sup>2</sup> | p | R<sup>2</sup> | p |
| Epiphytic | 2019 | 0.31 | **0.001** |  |  |  |  |
|  | 2020 | 0.26 | **0.0009** |  |  |  |  |
|  | 2021 | 0.23 | **0.0009** |  |  |  |  |
|  | All | 0.08 | **0.0009** | 0.2 | **0.0009** | 0.13 | **0.0009** |
| Endophytic | 2019 | 0.37 | **0.0009** |  |  |  |  |
|  | 2020 | 0.23 | **0.0009** |  |  |  |  |
|  | 2021 | 0.29 | **0.0009** |  |  |  |  |
|  | All | 0.1 | **0.0009** | 0.16 | **0.0009** | 0.11 | **0.0009** |

| **B** |  | R<sup>2</sup> | p |
| --- | --- | --- | --- |
| Combining all years,<br>landraces, and<br>communities | Landrace | 0.05 | **0.0009** |
|  | Year | 0.11 | **0.0009** |
|  | Community type | 0.06 | **0.0009** |
|  | Landrace x Year | 0.07 | **0.0009** |
|  | Landrace x Community type | 0.02 | **0.0009** |
|  | Year x community type | 0.04 | **0.0009** |
|  | Landrace x Year x community type | 0.036 | **0.0009** |

Finally, using a full model, we tested the effect of landrace, year, and community type on community composition, observing that sampling year explained the highest amount of variation in the dataset (11%), with weaker effects of landrace (5%) and community type (6%), and significant but weak interactions between variables (Table 1B). Together with the high variation in communities across years, these results suggest that both types of phyllosphere communities are more strongly structured by environmental factors rather than host identity, with relatively weak differential effects across epiphytic vs. endophytic communities. Nonetheless, the differences were generally in the predicted direction, with epiphytic communities being more variable.

### No evidence for vertical transmission of leaf phyllosphere communities

Next, we tested whether the relatively weak effect of host landrace on the phyllosphere microbiome could arise from inconsistent vertical transmission of microbial communities across generations. For this analysis, we used seeds of the landrace Chakhao (CK), finding that the seed epiphytic and endophytic communities both varied significantly across years, with a higher impact of sampling year on the epiphytes (epiphytes, PERMANOVA, seed_2019 vs 2020_, R^2^ = 0.34, p = 0.0009; endophytes, PERMANOVA, seed_2019 vs 2020,_ R^2^ = 0.18, p = 0.0009)

(Fig 5). In contrast, the leaf microbiomes were more stable, and many of the bacterial genera detected in the seed epiphytic or endophytic communities did not colonize the leaf epiphytic or endophytic niches in the next generation (Fig 5). Indeed, sampling year had a larger impact on community structure than plant niche or community type (PERMANOVA, effect of niche (leaf/seed): R^2^ = 0.08, p = 0.0009; effect of community type: R^2^ = 0.03, p = 0.0009; effect of year: R^2^ = 0.13, p = 0.0009; niche x community type: R^2^ = 0.02, p = 0.004; niche x year: R^2^ = 0.04, p = 0.0009; community type x year, R^2^ = 0.04, p = 0.0009, niche x community type x year: R^2^ = 0.015, p = 0.0009). Together, these results support the idea that weak vertical transmission contributes to stronger environmental effects on rice phyllosphere microbiomes.

**Fig 5:**
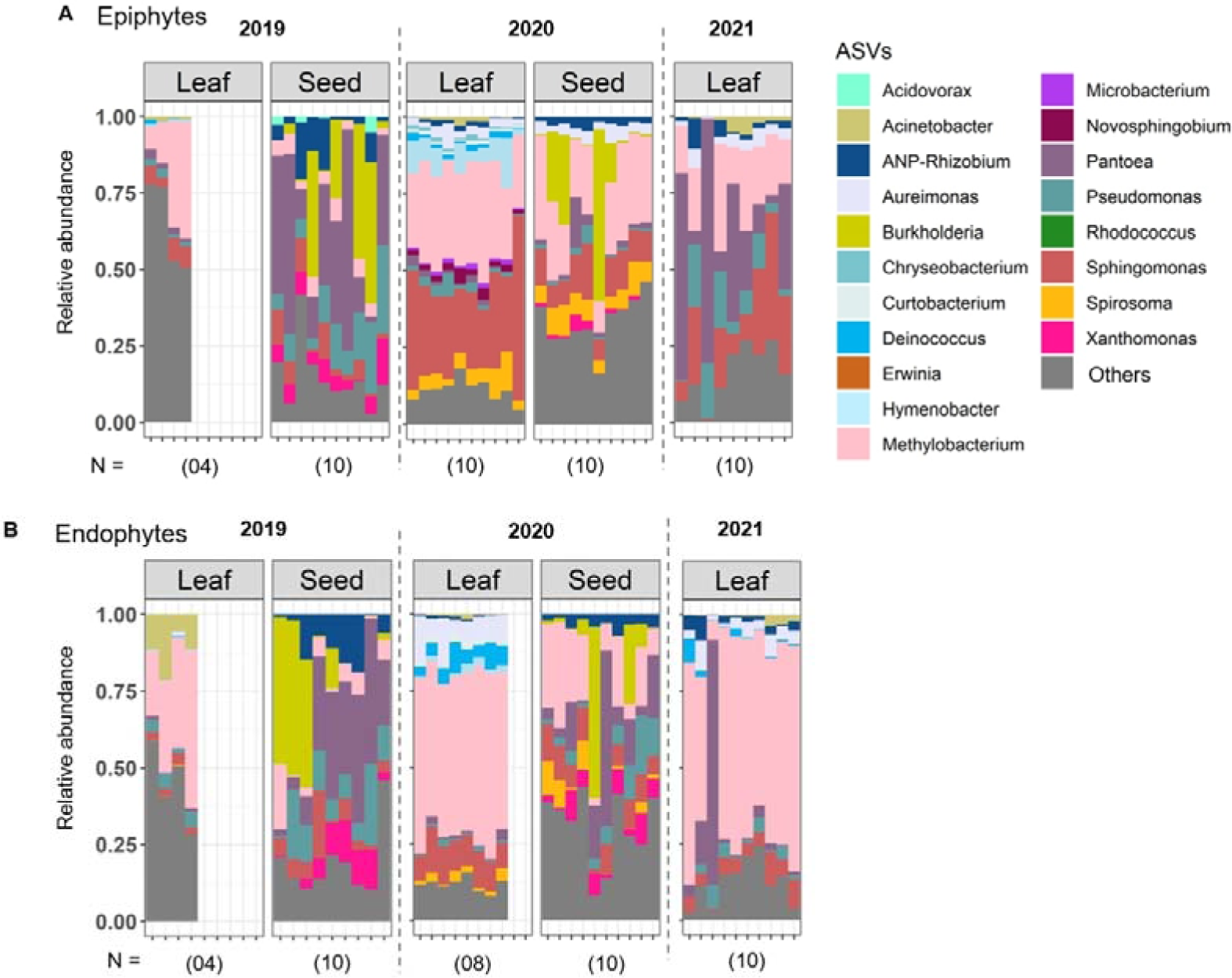
Leaf and seed communities vary across generations of Chakhao plants. Stacked bar plots show the relative abundance of bacterial ASVs (genus level) across the leaf and seeds of Chakhao as (A) epiphytes and (B) endophytes. Sample size (number of replicate plants) is indicated in parentheses. Taxa with <1% reads, or that were not assigned to known bacteria in the database, were combined into “others”.

### Epiphytic bacterial communities have higher modularity and co-occurrence networks with more positive correlations, indicative of lower stability

Lastly, to test our third prediction about the relative stability and consistency of epiphytic vs. endophytic communities, we analysed the network of co-occurrence of taxa. We estimated the correlation coefficients for the relative abundance of each pair of bacterial genera across landraces (in a given year) or across years (for a given landrace), retaining only significant correlation coefficients with absolute values greater than 0.5 (see Methods). We used these correlations to generate a co-occurrence network. Based on prior work (52), we expected that the more variable epiphytic communities would show higher modularity in the co-occurrence network, whereas we would see simpler, less modular networks for endophytic communities. Indeed, in both 2019 and 2020, epiphytic communities had more dense networks with more modules (Fig 6).

**Fig 6:**
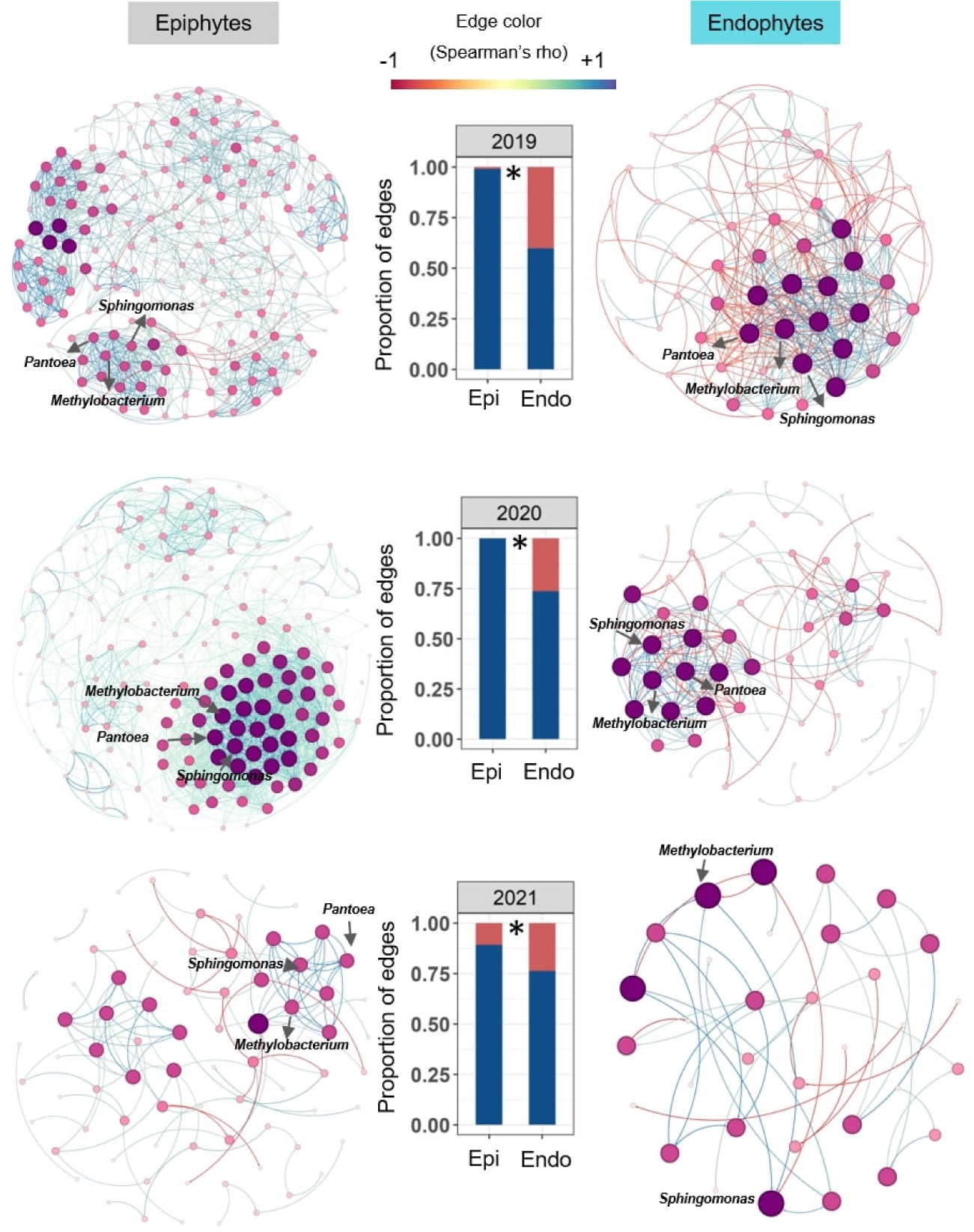
Epiphytic bacterial communities have higher modularity and more complex co-occurrence networks. Networks indicate co-occurrence relationships between bacterial genera across the three sampling years, with each node representing a genus. Networks were constructed using data pooled for all four focal landraces (see methods section for details). Node size and color both indicate the degree of connectivity, with larger node sizes and darker colors indicating nodes with more connections. The edge colors blue and red represent positive and negative relationships between nodes. Dominant and core bacterial genera are indicated in the networks. Bar plots show the proportion of pairwise relationships that are positive vs. negative (blue vs. red, respectively), in both types of communities (asterisks indicate significant differences, p<0.05, Chi square test).

Another important feature of these networks is the nature of the correlation between genera. Prior simulations suggest that a mixed set of correlations (i.e., some positive and some negative) increase network stability, compared to networks with only positive relationships (52). Epiphytic communities showed almost exclusively positive relationships among taxa in all three years, whereas endophytic communities were more balanced (Fig 6). This pattern held true even when we considered all landraces for a given year (Fig S8), or if we excluded the 2019 epiphytic community of Chakhao (CK), which had a very high proportion of rare taxa (Fig S9). The overall variability in the networks is exemplified by focusing on the genus *Methylobacterium*, the most dominant genus in our dataset, which was significantly correlated with different bacterial genera across years, in both types of communities (Fig S10). However, *Methylobacterium* did have strong and consistent positive associations with the genera *Aureimonas* and *Deinococcus* in endophytic and epiphytic communities respectively (Fig S10). Finally, we also compared co-occurrence networks across each of our four focal landraces. The networks varied substantially across landraces, but again showed high modularity with higher proportions of positive relationships in epiphytic microbiomes, except in the case of landrace Tolenphou (TP), where epiphytic and endophytic community structures were similar (Fig S11). Together, these results support our hypothesis that endophytic communities are more stable than phyllosphere epiphytic communities, across years as well as across host landraces.

### Divergence in phyllosphere communities arises during the flowering stage of rice

Overall, the results described above support our prediction that endophytic communities of rice landraces are less variable and more robust than epiphytic communities. Does this difference arise early in the plant’s life cycle, or emerge later during growth? To address this, in the last year of our fieldwork (2021), we collected leaf samples across different growth stages for one of the rice landraces in our study (Chakhao, CK), and analyzed microbiome changes as the plants developed. The epiphytic community composition differed significantly across growth stages, with several late-colonizing genera (e.g., *Muribacter, Rodentibacter, Rothia, Streptobacillus,* and *Streptococcus*) being lost or reduced in proportion as plants matured (Fig 7A). In contrast, the composition of the endophytic community was more consistent, with 70-80% of the microbiome composed of the genus *Methylobacterium* across growth stages (Fig 7A). Host plant growth stage had a slightly weaker effect in endophytic communities (effect of growth stage, endophytic community: R^2^ = 0.29, p < 0.001, epiphytic community: R^2^ = 0.23, p < 0.001) (Fig S13A), supported by a PERMANOVA analysis of full community composition showing a significant but weak growth stage x community type interaction (growth stage effect, R^2^ = 0.14, p < 0.001; community type, R^2^ = 0.14, p < 0.001; growth stage x community type, R^2^ = 0.07, p < 0.001).

**Fig 7:**
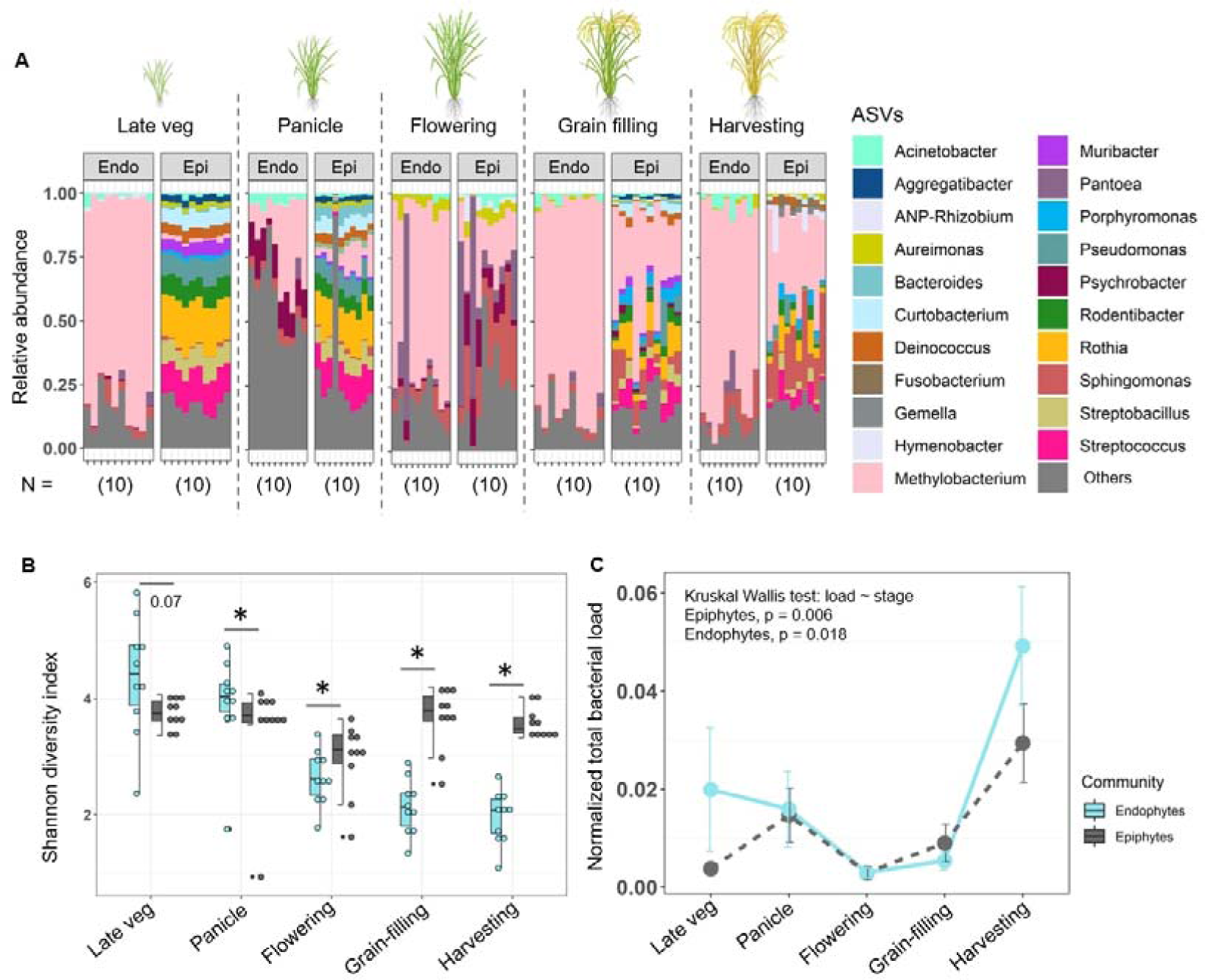
Phyllosphere microbiomes vary across growth stages of the host rice plant. (Chakhao): (A) Community composition of each plant sampled at different host growth stages. (B) Shannon diversity index of epiphytic vs. endophytic communities at different growth stages. n=10 replicate plants for each growth stage; asterisks represent significant pairwise differences between community types at the specific growth stage (paired t-test, p<0.05). (C) Total bacterial load in each plant sampled at specific growth stages (n=5), normalized to the total leaf tissue as measured from the endophytic sample (the same leaf sample was used to extract both community types, as described in the Methods). Error bars show standard error.

In the earlier stages of host growth (late vegetative and panicle formation), endophytic communities tended to have higher diversity and richness than epiphytes, but this pattern flipped at the flowering stage (pairwise tests shown in Fig 7B; Kruskal Wallis tests, richness ∼ stage (epiphytes), p = 0.0009; richness ∼ stage (endophytes), p = 6.3 X 10^-7^); diversity ∼ stage (epiphytes), p = 0.004; diversity ∼ stage (endophytes), p = 2.02 X 10^-6^) (Fig 7B, Fig S12). Thus, by the time we sampled microbiomes in the experiments described earlier (flowering stage), the diversity and richness patterns had already reversed and matched our predictions. Overall, the growth stage and its interaction with community type significantly influenced community richness (ANOVA, richness ∼ stage x community; p _stage_ = 4.18 X 10^-9^;p _community_ = 0.8; p _stage X community_ = 4.16 X 10^-5^), while all factors significantly affected community diversity (ANOVA, diversity ∼ stage X community; p _stage_ = 1.31 X 10^-10^;p _community_ = 0.000195; p _stage X community_ = 9.56 X 10^-9^). The decreasing diversity and richness of the endophytic community across growth stages suggests that many of the bacterial taxa are filtered out over time, potentially due to host selection.

We then asked whether the identified core bacterial taxa (shown in previous sections) were abundant throughout the growth stages or colonized at a specific growth stage of the rice plant. Many of the core bacterial genera (*Aureimonas, Methylobacterium, Pantoea, Pseudomonas, and Sphingomonas)* were most dominant at the flowering stage (Fig S13B). However, the total bacterial load in leaf samples (relative to leaf tissue) did not increase at the flowering stage, but instead peaked at the harvesting stage (Fig 7C). Reflecting our previous results for 2021 communities, the total load was not correlated with either diversity or richness of either type of community (Fig S14).

## DISCUSSION

Our field study with paired sampling of epiphytic and endophytic phyllosphere communities highlights key factors governing community assembly, and uncovers points of convergence as well as divergence in the microbiomes of the two distinct leaf niches. We observed broad convergence in patterns of variation of both types of communities across time as well as host landraces. Very low individual variation across plants within a landrace, combined with significant differences across landraces cultivated in the same field in close proximity, suggest that the landrace effect likely reflects deterministic variation. This effect may reflect local adaptation to host-imposed divergent selection, e.g., as a function of host genotypes (e.g., see (53), and/or mediated by differences in host phenotypes such as leaf physical traits (35), nutrient availability (54), or production of hormones such as cytokinins (55) and ethylene (6). Further work is required to test these possibilities, and to quantify the role of stochastic vs. deterministic processes.

Despite the broad similarities in factors governing microbiome community composition and core community members, we observed several points of divergence between communities inhabiting the two leaf niches, beginning with distinct community composition. Epiphytic communities have higher richness and diversity, and are more variable across individual plants, as expected given their greater exposure to the environment. Interestingly, epiphytic communities also have higher bacterial loads. A network analysis revealed more modules in epiphytic communities, with greater positive associations between taxa. Previous work suggests that such communities, with more positive associations between members, are temporally more unstable (52). Our results therefore add to the growing body of work comparing epiphytic vs. endophytic communities of different host plants, suggesting that their distinct roles and consequences (discussed in the Introduction) may be attributed to differences in community composition, stability, and individual variation.

Another important difference across epiphytic and endophytic communities is the relative impact of environment vs. host. As expected, focusing on our four main landraces we find a stronger impact of sampling year on the epiphytic than the endophytic community. We also expected a stronger impact of host rice landrace and more predictable community members in the endophytic community. However, the variation explained by host rice landrace is only slightly higher in endophytic communities, and the landrace effect is weaker than the effect of sampling year on both communities, even when combining data from all landraces and years. Why do we observe a stronger effect of sampling year compared to host landrace? One possibility is that temporal variation in abiotic conditions was extremely high in our sampling years, overwhelming the baseline effects of host genotype. Another cause may be the lack of vertical transmission of phyllosphere communities across generations via seeds, in turn leading to a larger impact of environmental factors (which likely vary across years). Third, despite large phenotypic differences, our host landraces may be physiologically very similar, and may therefore impose similar filters on their microbiomes. We find some support for this speculation: when we included two landraces with very distinct microbiomes in our analysis (MP and MR), the impact of host landrace increased (compare landrace effect in Table 1 vs. Table S2; Fig S3). Finally, our microbiome analysis is relatively coarse-grained, and prior work shows substantial hidden diversity across strains of a given species (56). Thus, more detailed taxonomic and genomic analysis of the dominant microbial taxa may reveal stronger host landrace effects if we could distinguish between functionally different strains or niche-specialist species. Further analysis is necessary to distinguish between these possibilities.

We also observed distinct patterns of variation in the two types of phyllosphere communities across plant growth stages. Focusing on Chakhao rice, we find that growth stage has a higher impact on the endophytic than epiphytic community, with the latter showing a more consistent richness and diversity over time. Similar results were previously reported in maize plants (57). Prior work with *Arabidopsis thaliana* also suggests greater stochasticity and dispersal limitation in epiphytic communities at the early stages and later deterministic host selection in microbiomes (58, 59). Thus, our results perhaps reflect broader patterns across plant species. Microbiome variation across growth stages may be driven by developmental shifts in host secondary metabolite profiles (60), or priority effects, where the presence of specific taxa at early stages alters subsequent community composition. For example, inoculating rice seeds with the endophyte *Xanthomonas sacchari* altered the stem and rhizosphere endophytic community significantly at later stages (61). Such priority effects may be especially important in leaf endophytic phyllosphere communities, leaving leaf epiphytic communities more prone to the effects of immigration.

Finally, the total bacterial loads in epiphytic and endophytic niches also tended to vary, with significantly higher loads in epiphytic communities of two landraces (Fig 3C). These results mirror prior work with *Arabidopsis* showing that CFUs (colony forming units) in epiphytes are ∼100-fold higher in the endophytic community (4). The total microbial load for both community types also varied significantly across landraces, potentially arising due to differences in leaf architecture or metabolic profiles across landraces. For instance, *Salmonella enterica* colonization varies with the abundance of type 1 trichomes on tomato leaves (62). Strikingly, the total microbial load in our study was not correlated with the microbiome diversity or richness for either community type, and this was also true across plant developmental stages. This was surprising because we generally expect that colonization by highly beneficial core taxa should increase microbial load, driving a negative relationship between diversity and load. We speculate that the lack of such a relationship in our samples reflects shared functional traits across several microbes, such that high colonization rate may be achieved by any of a set of taxonomically distinct but functionally similar bacteria. Another possibility is that plants may select for groups of interacting bacteria rather than single species or strains, allowing for high diversity to be maintained along with high microbial loads. This possibility is consistent with the strongly modular network architecture observed in our dataset. Yet another possibility is that high microbial loads reflect transient pathogen infection rather than colonization by commensals or symbionts. However, this is unlikely in our case because we did not find known pathogens such as *Xanthomonas oryzae* in any of our communities. Finally, we note that for logistical simplicity we used the 16S rRNA gene to measure absolute bacterial load, but this is not ideal because of copy number variation in this gene across bacteria. Further studies with single-copy genes would provide more accurate estimates of total bacterial load (63, 64).

Supporting our prior finding that *Methylobacterium* can be beneficial for rice hosts (33) we find the genus *Methylobacterium* to be one of the most abundant and consistent core bacteria for both phyllosphere community types in all landraces. *Methylobacterium* is a widespread plant-associated bacterial genus with the ability to utilize methanol released by plants as a carbon source, in turn promoting plant growth via several mechanisms (cite a review here, we don’t need all the details in the next few sentences). However, the strength of the association between the host plant and *Methylobacterium* is variable (33, 65). This variability could potentially arise if several community members together influence host plant fitness. For instance, apart from *Methylobacterium* we identified several genera such as *Sphingomonas, Pantoea, Aureimonas*, and *Pseudomonas* as core bacterial taxa in both the phyllosphere community types — similar to a recent study with wild and cultivated rice (51) — some of which (*Sphingomonas* and *Pantoea*) are also known to promote plant growth (26, 66). We observed strong positive and consistent co-occurrence of *Methylobacterium* with *Aureimonas* and *Deinococcus* across years, landraces, and community types (Fig S10). We speculate that these correlations may indicate cooperative interactions that need experimental validation. Interestingly, nearly all of these genera are also consistent seed endophytes across generations, suggesting that co-inoculating rice with several co-occuring core taxa may be a fruitful avenue for further research aimed at increasing rice yield or growth.

In summary, our work identifies and quantifies several factors that shape the rice phyllosphere microbiome, finding points of both convergence and divergence in the assembly and maintenance of bacterial communities in two distinct leaf niches. We hope that further studies determine if and how the identified core bacterial taxa impact their host rice landrace, and investigate potential agricultural applications.

## ACKNOWLEDGEMENTS

We thank W. Ibobo Singh and Sharat, KVK Thoubal for allowing us access to their fields for sampling; S Chandrakanta Singh for sample collection in 2020, during the COVID-19 pandemic; and Nitish Malhotra and Rittik Deb for help with R scripts. We acknowledge funding and support from the Department of Biotechnology (grant DBT – 358 NER/AGRI/24/2013), the National Centre for Biological Sciences (NCBS–TIFR) and the Department of Atomic Energy, Government of India (Project Identification No. RTI 4006).

## AUTHOR CONTRIBUTIONS

PS conceived, designed, conducted experiments, analysed data, and drafted the manuscript. DA conceived and designed experiments, acquired funding, and wrote the manuscript.

## DATA AVAILABILITY

Raw sequence data are available in the Sequence Read Archive (SRA) of the National Centre for Biotechnology Information (NCBI) with SRA accession numbers (X) under Bioproject (X). All other data supporting the findings are included as Supplementary file SI_Datasheet1.

## REFERENCES

1. Vorholt JA. 2012. Microbial life in the phyllosphere. Nat Rev Microbiol 10:828–840.

2. Vacher C, Hampe A, Porté AJ, Sauer U, Compant S, Morris CE. 2016. The Phyllosphere: Microbial Jungle at the Plant-Climate Interface. Annu Rev Ecol Evol Syst 47:1–24.

3. Bashir I, War AF, Rafiq I, Reshi ZA, Rashid I, Shouche YS. 2022. Phyllosphere microbiome: Diversity and functions. Microbiol Res 254.

4. Chen T, Nomura K, Wang X, Sohrabi R, Xu J, Yao L, Paasch BC, Ma L, Kremer J, Cheng Y, Zhang L, Wang N, Wang E, Xin XF, He SY. 2020. A plant genetic network for preventing dysbiosis in the phyllosphere. Nature 580:653–657.

5. Fitzpatrick CR, Salas-González I, Conway JM, Finkel OM, Gilbert S, Russ D, Teixeira PJPL, Dangl JL. 2020. The Plant Microbiome: From Ecology to Reductionism and beyond. Annu Rev Microbiol 74:81–100.

6. Bodenhausen N, Bortfeld-miller M, Ackermann M, Vorholt JA. 2014. A Synthetic Community Approach Reveals Plant Genotypes Affecting the Phyllosphere Microbiota 10.

7. Wagner MR, Lundberg DS, Rio TG, Tringe SG, Dangl JL, Mitchell-olds T. 2016. Host genotype and age shape the leaf and root microbiomes of a wild perennial plant. Nat Commun 7:1–15.

8. Schlechter RO, Miebach M, Remus-emsermann MNP. 2019. Driving factors of epiphytic bacterial communitiesLJ: A review. J Adv Res 19:57–65.

9. Zhu YG, Xiong C, Wei Z, Chen QL, Ma B, Zhou SYD, Tan J, Zhang LM, Cui HL, Duan GL. 2022. Impacts of global change on the phyllosphere microbiome. New Phytol 234:1977–1986.

10. Huang S, Zha X, Fu G. 2023. Affecting Factors of Plant Phyllosphere Microbial Community and Their Responses to Climatic Warming—A Review. Plants 12:1–11.

11. Firrincieli A, Khorasani M, Frank AC, Doty SL. 2020. Influences of Climate on Phyllosphere Endophytic Bacterial Communities of Wild Poplar. Front Plant Sci 11:1– 13.

12. Gomes T, Pereira JA, Benhadi J, Lino-Neto T, Baptista P. 2018. Endophytic and Epiphytic Phyllosphere Fungal Communities Are Shaped by Different Environmental Factors in a Mediterranean Ecosystem. Microb Ecol 76:668–679.

13. Al Ashhab A, Meshner S, Alexander-Shani R, Dimerets H, Brandwein M, Bar-Lavan Y, Winters G. 2021. Temporal and Spatial Changes in Phyllosphere Microbiome of Acacia Trees Growing in Arid Environments. Front Microbiol 12:1–14.

14. Morella NM, Weng FCH, Joubert PM, Jessica C, Lindow S, Koskella B. 2020. Successive passaging of a plant-associated microbiome reveals robust habitat and host genotype-dependent selection. Proc Natl Acad Sci U S A 117:1148–1159.

15. Redford AJ, Fierer N. 2009. Bacterial Succession on the Leaf SurfaceLJ: A Novel System for Studying Successional Dynamics 189–198.

16. Bao L, Gu L, Sun B, Cai W, Zhang S, Zhuang G, Bai Z, Zhuang X. 2019. Seasonal variation of epiphytic bacteria in the phyllosphere of Gingko biloba, Pinus bungeana and Sabina chinensis. FEMS Microbiol Ecol 96:1–13.

17. Massoni J, Bortfeld-Miller M, Jardillier L, Salazar G, Sunagawa S, Vorholt JA. 2020. Consistent host and organ occupancy of phyllosphere bacteria in a community of wild herbaceous plant species. ISME J 14:245–258.

18. Trivedi P, Leach JE, Tringe SG, Sa T, Singh BK. 2020. Plant–microbiome interactions: from community assembly to plant health. Nat Rev Microbiol 18:607– 621.

19. Yu X, Lund SP, Scott RA, Greenwald JW, Records AH, Nettleton D, Lindow SE, Gross DC, Beattie GA. 2013. Transcriptional responses of Pseudomonas syringae to growth in epiphytic versus apoplastic leaf sites. Proc Natl Acad Sci U S A 110.

20. Meyer KM, Muscettola IE, Vasconcelos ALS, Sherman JK, Metcalf CJE, Lindow SE, Koskella B. 2023. Conspecific versus heterospecific transmission shapes host specialization of the phyllosphere microbiome. Cell Host Microbe 31:2067–2079.e5.

21. Wang P, Kong X, Chen H, Xiao Y, Liu H, Li X, Zhang Z, Tan X, Wang D, Jin D, Deng Y, Cernava T. 2021. Exploration of Intrinsic Microbial Community Modulators in the Rice Endosphere Indicates a Key Role of Distinct Bacterial Taxa Across Different Cultivars. Front Microbiol 12:1–12.

22. Demarquest G, Lajoie G. 2023. Bacterial endophytes of sugar maple leaves vary more idiosyncratically than epiphytes across a large geographic area. FEMS Microbiol Ecol 99:1–11.

23. Ali S, Ganai BA, Kamili AN, Bhat AA, Mir ZA, Bhat JA, Tyagi A, Islam ST, Mushtaq M, Yadav P, Rawat S, Grover A. 2018. Pathogenesis-related proteins and peptides as promising tools for engineering plants with multiple stress tolerance. Microbiol Res 212–213:29–37.

24. Delmotte, NathanaelKnief C, Chaffron S, Innerebner G, Roschitzki B, Schlapbach R. 2009. Community proteogenomics reveals insights into the physiology of phyllosphere bacteria. Proc Natl Acad Sci 106:16428–16433.

25. Kumar M, Kour D, Yadav AN, Saxena R, Rai PK, Jyoti A, Tomar RS. 2019. Biodiversity of methylotrophic microbial communities and their potential role in mitigation of abiotic stresses in plants. Biologia (Bratisl) 74:287–308.

26. Asaf S, Numan M, Khan AL, Al-Harrasi A. 2020. Sphingomonas: from diversity and genomics to functional role in environmental remediation and plant growth. Crit Rev Asaf S, Numan M, Khan AL, Al-Harrasi A 2020 Sphingomonas from Divers genomics to Funct role Environ Remediat plant growth Crit Rev Biotechnol 40138–152 Biotechnol 40:138–152.

27. Meena KK, Kumar M, Kalyuzhnaya MG. 2012. Epiphytic pink-pigmented methylotrophic bacteria enhance germination and seedling growth of wheat (*Triticum aestivum*) by producing phytohormone. Antonie Van Leeuwenhoek 101:777–786.

28. Tsavkelova EA, Egorova MA, Leontieva MR, Malakho SG, Kolomeitseva GL, Netrusov AI. 2016. Dendrobium nobile Lindl. seed germination in co-cultures with diverse associated bacteria. Plant Growth Regul 80:79–91.

29. Khan AL, Waqas M, Kang SM, Al-Harrasi A, Hussain J, Al-Rawahi A, Al-Khiziri S, Ullah I, Ali L, Jung HY, Lee IJ. 2014. Bacterial endophyte Sphingomonas sp. LK11 produces gibberellins and IAA and promotes tomato plant growth. J Microbiol 52:689–695.

30. Palberg D, Kisiała A, Jorge GL, Emery RJN. 2022. A survey of Methylobacterium species and strains reveals widespread production and varying profiles of cytokinin phytohormones. BMC Microbiol 22:1–17.

31. Knief C, Chaffron S, Stark M, Innerebner G. 2012. Metaproteogenomic analysis of microbial communities in the phyllosphere and rhizosphere of rice. ISME 6:1378– 1390.

32. Matsumoto H, Fan X, Wang Y, Kusstatscher P, Duan J, Wu S, Chen S, Qiao K, Wang Y, Ma B, Zhu G, Hashidoko Y, Berg G, Cernava T, Wang M. 2021. Bacterial seed endophyte shapes disease resistance in rice. Nat Plants 7:60–72.

33. Sanjenbam P, Shivaprasad P V., Agashe D. 2022. Impact of Phyllosphere Methylobacterium on Host Rice Landraces. Microbiol Spectr 10.

34. Berg M, Koskella B. 2018. Nutrient- and Dose-Dependent Microbiome-Mediated Protection against a Plant Pathogen. Curr Biol 28:2487–2492.e3.

35. Leveau JH. 2019. A brief from the leaf: latest research to inform our understanding of the phyllosphere microbiome. Curr Opin Microbiol 49:41–49.

36. Medhabati K, Rajiv Das K, Rohinikumar M, Sunitibala H, Dikash Singh T. 2013. Genetic Divergence in Indigenous Wild and Cultivated Rice Species of Manipur Valley. ISRN Genet 2013:1–6.

37. Das B, Sengupta S, Parida SK, Roy B, Ghosh M, Prasad M. 2013. Genetic diversity and population structure of rice landraces from Eastern and North Eastern States of India. BMC Genet 14:1–14.

38. Roy S, Banerjee A, Mawkhlieng B, Misra AK. 2015. Genetic Diversity and Population Structure in Aromatic and Quality Rice (*Oryza sativa L.*) Landraces from North-Eastern India Somnath. PLoS One 10:1–13.

39. Umakanth B, Vishalakshi B, Sathish Kumar P, Rama Devi SJS, Bhadana VP, Senguttuvel P, Kumar S, Sharma SK, Sharma PK, Prasad MS, Madhav MS. 2017. Diverse Rice Landraces of North-East India Enables the Identification of Novel Genetic Resources for Magnaporthe Resistance. Front Plant Sci 8:1–13.

40. Fadrosh DW, Bing Ma PG, Sengamalay N, Ott S, Brotman RM, Ravel J. 2014. An improved dual-indexing approach for multiplexed 16S rRNA gene sequencing on the Illumina MiSeq platform. Microbiome 2:1–7.

41. Lundberg DS, Yourstone S, Mieczkowski P, Jones CD, Dangl JL. 2013. Practical innovations for high-throughput amplicon sequencing. Nat Methods 10:999–1002.

42. CoreTeam R. 2016. R Core Team. R: A Language and Environment for Statistical Computing. Vienna, Austria: R Foundation for Statistical Computing.

43. Callahan BJ, McMurdie PJ, Rosen MJ, Han AW, Johnson AJA, Holmes SP. 2016. DADA2: High-resolution sample inference from Illumina amplicon data. Nat Methods 13:581–583.

44. Oksanen J, Blanchet FG, Friendly M, Kindt R, Mcglinn D, Minchin PR, Hara RBO, Simpson GL. 2019. vegan: Community Ecology Package.R package version 2.5–4. https://CRAN.R-project.org/package=vegan.

45. Anderson MJ, Willis TJ. 2003. Canonical analysis of principal coordinates: A useful method of constrained ordination for ecology. Ecology 84:511–525.

46. Kindt R, Coe R. 2005. Tree diversity analysis.

47. Wickham H. 2016. ggplot2: Elegant Graphics for Data Analysis. Springer-VerlagNew York.

48. ning-et-al-2019-a-general-framework-for-quantitatively-assessing-ecological-stochasticity.pdf.crdownload.

49. Chase JM, Kraft NJB, Smith KG, Vellend M, Inouye BD. 2011. Using null models to disentangle variation in community dissimilarity from variation in α-diversity. Ecosphere 2.

50. Bastian M, Heymann S, Jacomy M. 2009. Gephi: An Open Source Software for Exploring and Manipulating Networks Visualization and Exploration of Large Graphs. Proc Thrid Int ICWSM Conf 361–362.

51. Yin Y, Wang Y-F, Cui H-L, Zhou R, Li L, Duan G-L, Zhu Y-G. 2023. Distinctive structure and assembly of phyllosphere microbial. Microbiol Spectr 11:1–14.

52. Coyte KZ, Schluter J, Foster KR. 2015. The ecology of the microbiome: Networks, competition, and stability. Science (80-) 350:663–666.

53. Petipas RH, Geber MA, Lau JA. 2021. Microbe-mediated adaptation in plants. Ecol Lett 24:1302–1317.

54. Thapa S, Prasanna R, Ranjan K, Velmourougane K. 2017. Nutrients and host attributes modulate the abundance and functional traits of phyllosphere microbiome in rice. Microbiol Res 204:55–64.

55. Gupta R, Elkabetz D, Leibman-Markus M, Sayas T, Schneider A, Jami E, Kleiman M, Bar M. 2022. Cytokinin drives assembly of the phyllosphere microbiome and promotes disease resistance through structural and chemical cues. ISME J 16:122– 137.

56. Karasov TL, Almario J, Friedemann C, Ding W, Giolai M, Heavens D, Kersten S, Lundberg DS, Neumann M, Regalado J, Neher RA, Kemen E, Weigel D. 2018. Arabidopsis thaliana and Pseudomonas Pathogens Exhibit Stable Associations over Evolutionary Timescales. Cell Host Microbe 24:168–179.e4.

57. Xiong C, Singh BK, He JZ, Han YL, Li PP, Wan LH, Meng GZ, Liu SY, Wang JT, Wu CF, Ge AH, Zhang LM. 2021. Plant developmental stage drives the differentiation in ecological role of the maize microbiome. Microbiome 9:1–15.

58. Beilsmith K, Perisin M, Bergelson J. 2021. Natural bacterial assemblages in arabidopsis thaliana tissues become more distinguishable and diverse during host development. MBio 12:1–16.

59. Maignien L, DeForce EA, Chafee ME, Murat Eren A, Simmons SL. 2014. Ecological succession and stochastic variation in the assembly of Arabidopsis thaliana phyllosphere communities. MBio 5:1–10.

60. Pontonio E, Di Cagno R, Tarraf W, Filannino P, De Mastro G, Gobbetti M. 2018. Dynamic and assembly of epiphyte and endophyte lactic acid bacteria during the life cycle of Origanum vulgare L. Front Microbiol 9:1–16.

61. Wang X, He S, He Q, Ju Z, Ma Y, Wang Z, Han J, Zhang X, Wang X, He S, He Q, Ju Z, Ma Y, Wang Z, Han J, Zhang X. 2023. Bacterial Communities Across Rice Plant Growth Stages. Microbiol Spectr 0.

62. Barak JD, Kramer LC, Hao LY. 2011. Colonization of tomato plants by salmonella enterica is cultivar dependent, and type trichomes are preferred colonization sites. Appl Environ Microbiol 77:498–504.

63. Case RJ, Boucher Y, Dahllöf I, Holmström C, Doolittle WF, Kjelleberg S. 2007. Use of 16S rRNA and rpoB genes as molecular markers for microbial ecology studies. Appl Environ Microbiol 73:278–288.

64. Ogier JC, Pagès S, Galan M, Barret M, Gaudriault S. 2019. RpoB, a promising marker for analyzing the diversity of bacterial communities by amplicon sequencing. BMC Microbiol 19:1–16.

65. Tani A, Sahin N, Fujitani Y, Kato A, Sato K, Kimbara K. 2015. *Methylobacterium* species promoting rice and barley growth and interaction specificity revealed with whole-cell matrix-assisted laser desorption / ionization-timeof-flight mass spectrometry (MALDI-TOF/ MS) analysis. PLoS One 10:1–15.

66. Lv L, Luo J, Ahmed T, Zaki HEM, Tian Y, Shahid MS, Chen J, Li B. 2022. Rice Production 1–21.

