## Supplementary tables and figures for "Divergence and convergence in epiphytic and endophytic phyllosphere bacterial communities of rice landraces"

**Table S1: List of rice landraces and year of leaf sample collection**. Sample sizes indicate the number of individual plants sampled (1 leaf section per plant). Four focal rice landraces and one commercial variety (HYV) were consistently sampled and are described in the main text, and these are highlighted in bold.

| **Landraces** | **Sample size** | | | **Field sites** |
| --- | --- | --- | --- | --- |
|  | **2019** | **2020** | **2021** |  |
| **Chakhao (CK)** | **5** | **10** | **10** | Chingarel |
| **Phouren-mubi (PM)** | **5** | **10** | **10** | Chingarel |
| **Phoungang (PN)** | **5** | **10** | **10** | Andro |
| **Tolenphou (TP)** | **5** | **10** | **10** | Andro |
| **High Yielding variety (HYV)** | **5** | **10** | **10** | Chingarel |
| Chakhao khongnambi (CKS) | 5 | - | - | KVK Thoubal |
| Chakhao (CKA) | - | - | 5 | Andro |
| Kakcheng phou (KAK) | 5 | - | - | KVK Thoubal |
| Kumbi phou (KUM) | 5 | - | - | KVK Thoubal |
| Moirangphou kvk (KHOK) | 5 | - | - | KVK Thoubal |
| Moirangphou (MP) | 5 | 7 | - | Andro |
| Moirangphou khokngangbi (MR) | - | 7 | 10 | Andro |
| Phouren (PKVK) | 5 | - | - | KVK Thoubal |

**Table S2: Full model testing using data for all landraces.** Effect of landrace, and year on both epiphytic and endophytic phyllosphere microbial community for all rice varieties listed in Table S1, testing using separate PERMANOVAs. Significant P-values are shown in bold.

|  |  | Landrace | | Year | | Landrace X Year | |
| --- | --- | --- | --- | --- | --- | --- | --- |
| Community | Year | R^2^ | P | R^2^ | P | R^2^ | P |
| Epiphytic | 2019 | 0.38 | **0.0009** |  |  |  |  |
|  | 2020 | 0.29 | **0.0009** |  |  |  |  |
|  | 2021 | 0.22 | **0.0009** |  |  |  |  |
|  | All | 0.12 | **0.0009** | 0.13 | **0.0009** | 0.12 | **0.0009** |
| Endophytic | 2019 | 0.45 | **0.0009** |  |  |  |  |
|  | 2020 | 0.29 | **0.0009** |  |  |  |  |
|  | 2021 | 0.31 | **0.0009** |  |  |  |  |
|  | All | 0.15 | **0.0009** | 0.09 | **0.0009** |  |  |

**SUPPLEMENTARY FIGURES**

**Fig S1**: **Experimental design**. Schematic summarizing (A) the sampling year and number of landraces collected for DNA extraction (B) samples collected across growth stages of landrace Chakhao.

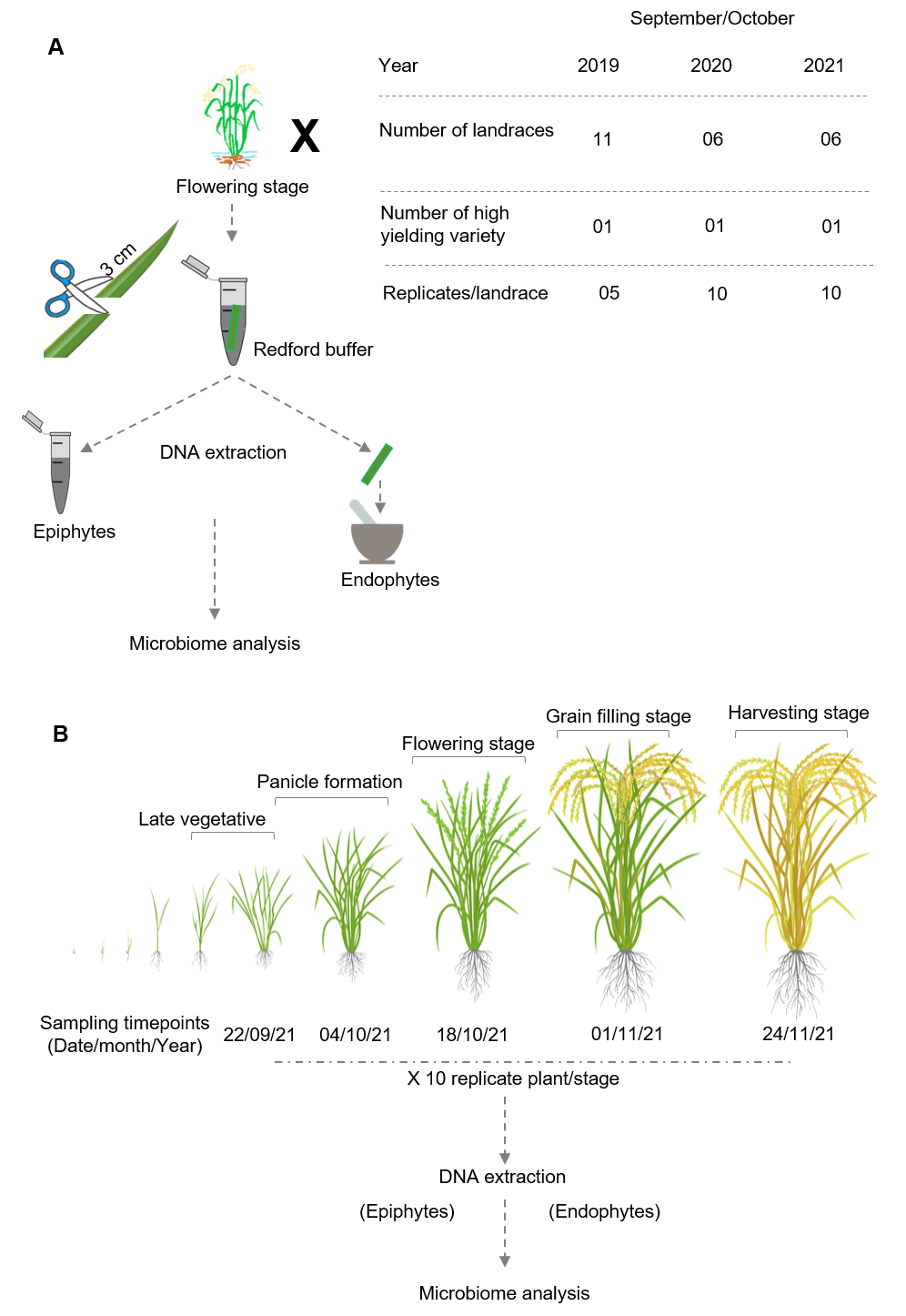

**Fig S2**: **Sequencing reads obtained per sample across years**. Total bacterial reads obtained from (A) epiphytic and (B) endophytic samples across three years. (C) Rarefaction curves showing the number of ASVs (richness) as a function of the number of reads sampled for the two communities. Each line represents a sample (sample names include landrace, year, and replicate; “E” indicates endophytes), and error bars indicate the standard error of richness estimated from 1000 random subsamples drawn from the total reads for each dataset. Dotted grey lines represent our threshold for the minimum number of total reads per sample required to proceed with further analysis.

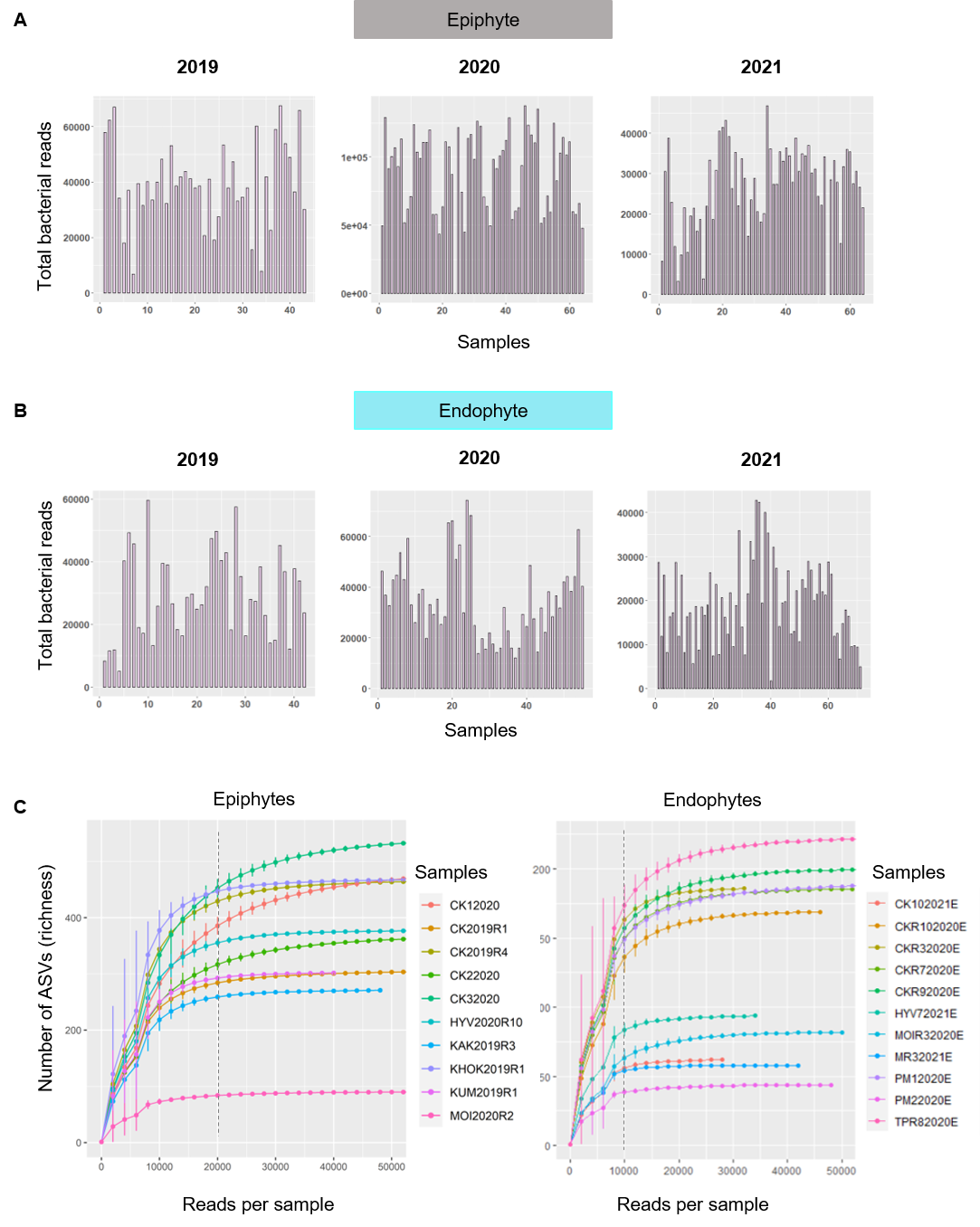

**Fig S3: Individual variation in host microbiomes for non-focal landraces**. Stacked bar plots showing the relative abundance of bacterial ASVs (identified to genus level) for (A) high yielding variety (HYV) (B) Moirangphou (MP) (C) Moirangphou khokngangbi (MR) (D) five landraces that were only sampled in 2019. Sample size (number of replicate plants) is indicated in parentheses. Reads less than 1% and unidentified taxa are clubbed as “others”.

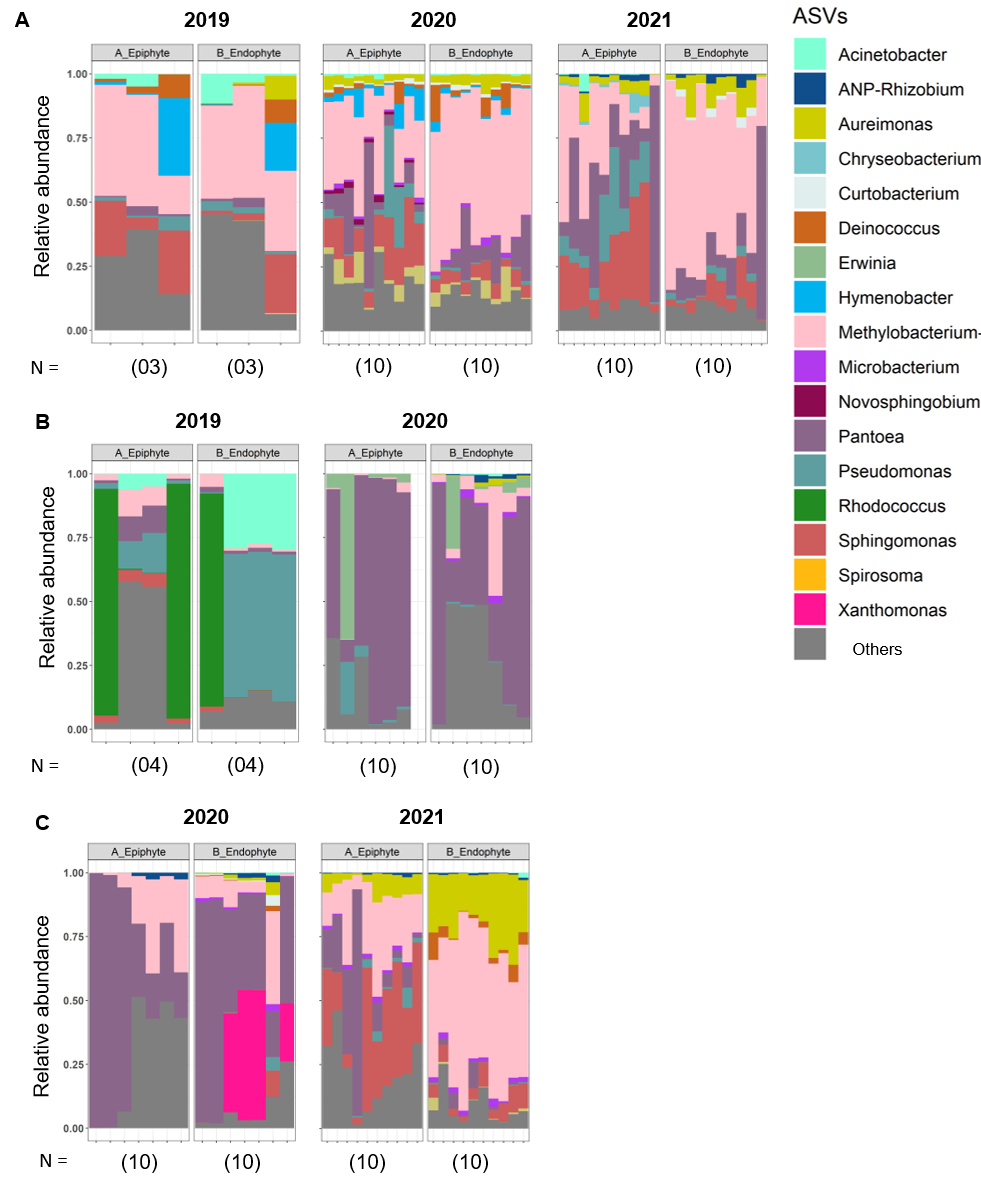

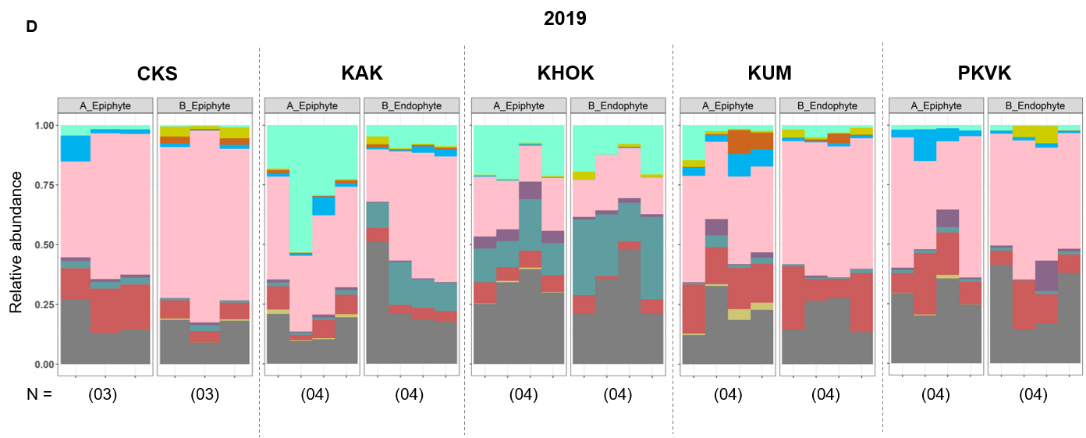

**Fig S4: Core *Methylobacterium* ASVs associated with rice landraces.** Heatmap showing the relative abundance of *Methylobacterium* ASVs (fraction of the total *Methylobacterium* reads) detected in >=10% of all samples. Core and dominant ASVs shared across both community types across years and landraces are highlighted by boxes.

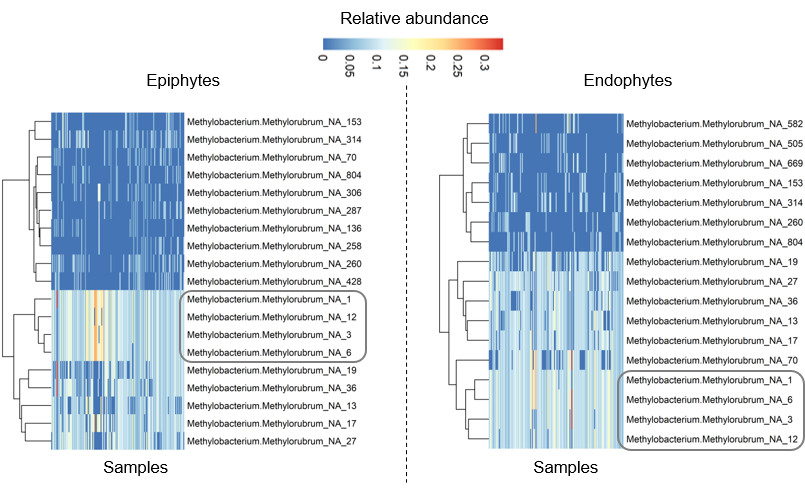

**Fig S5**: **Diversity and richness of** **microbial communities in non-focal landraces.** Boxplots show the Shannon diversity index and number of ASVs (richness) of both bacterial communities of all non-focal rice landraces sampled in a given year (paired t-test, P<0.05). Sample sizes are given in Table S1.

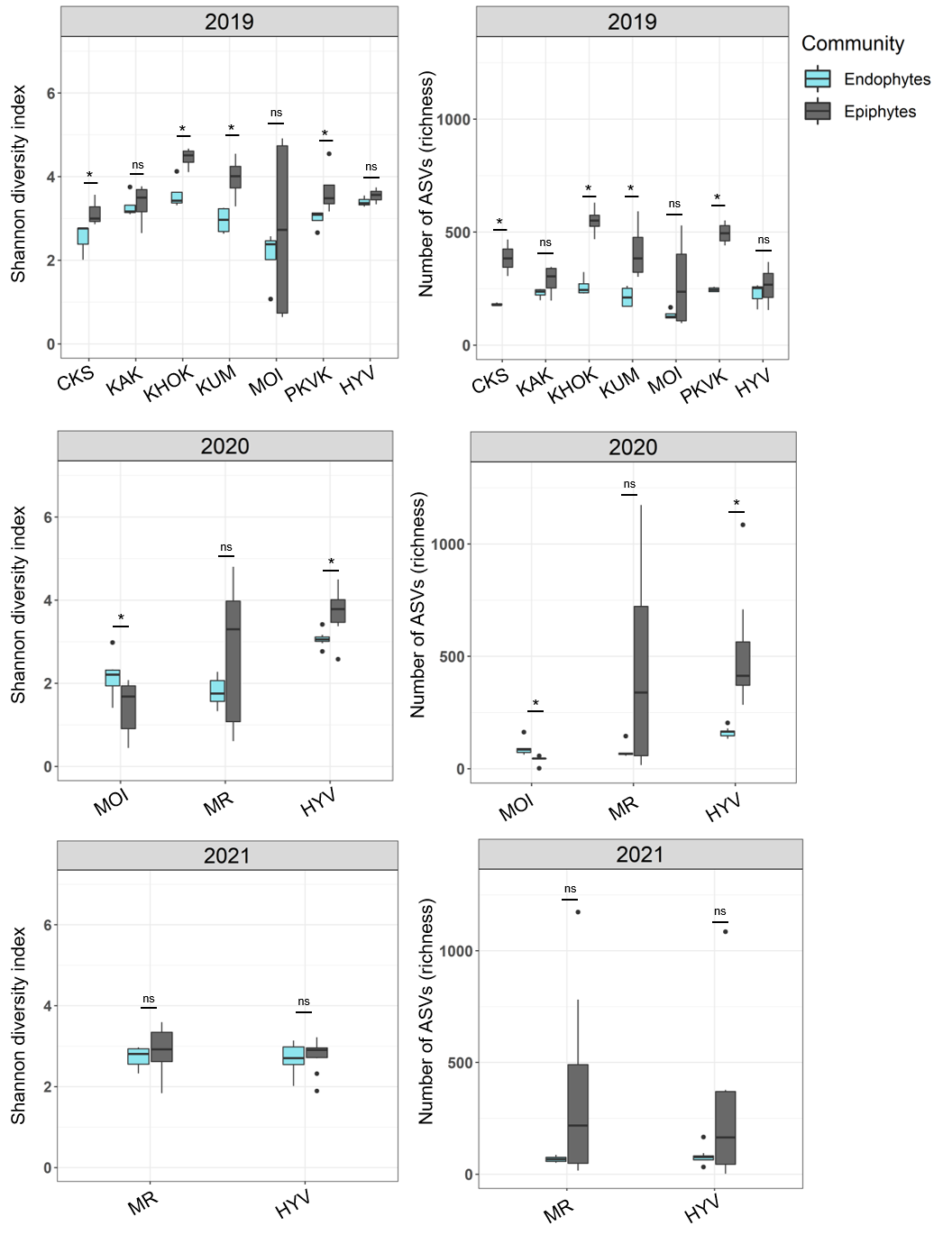

**Fig S6: Total bacterial load does not correlate with community diversity or richness**. Scatterplots showing the relationship between the normalized total bacterial load and Shannon diversity or richness across samples. Each point represents the mean value for 5 biological replicate plants of 4 rice landraces sampled in 2021. The output of non-parametric correlation tests is given in each panel.

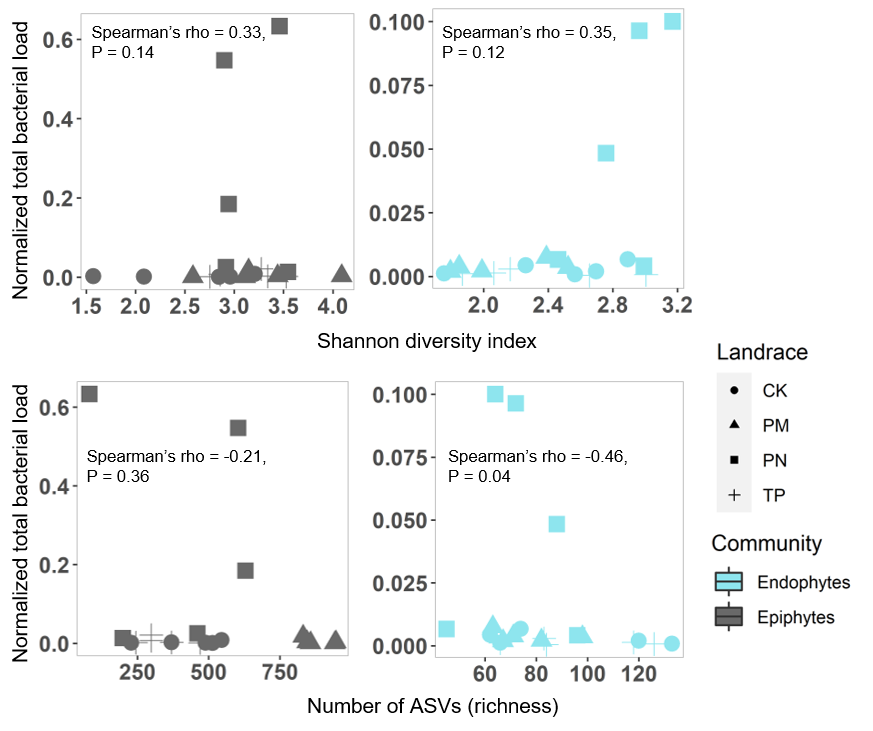

**Fig S7**: **Differentiation in microbial communities across host rice landraces**. Linear discriminant (LD) plots showing the clustering of (A) epiphytic and (B) endophytic bacterial communities across landraces and years. Numbers in parentheses indicate the number of biological replicates in each rice landrace. Axis labels indicate the proportion of variation explained, and ellipsoids represent 95% confidence intervals.

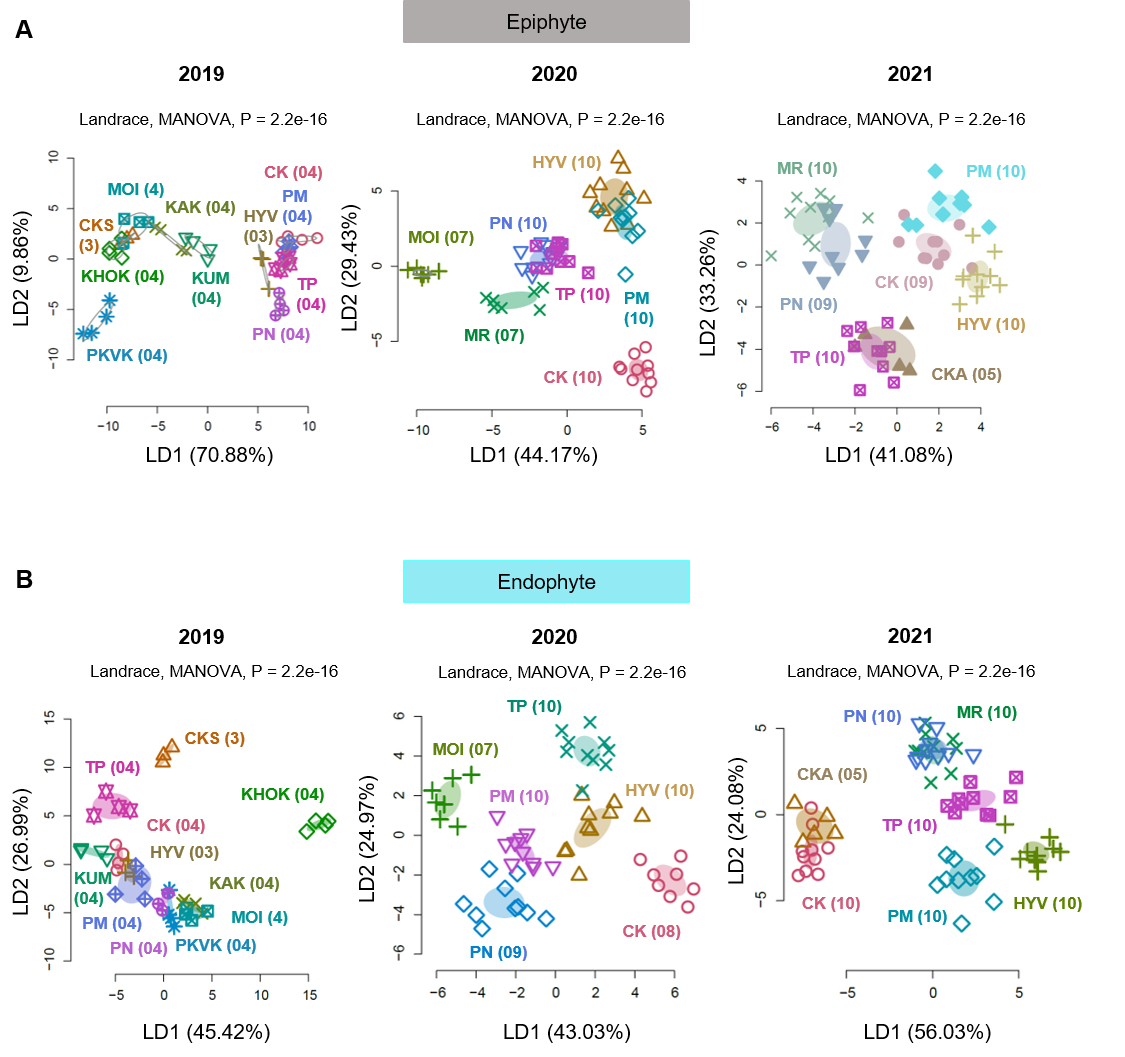

**Fig S8: Co-occurrence networks of epiphytic and endophytic bacterial communities across years.** Nodes represent a bacterial taxon (identified to genus level); larger and darker nodes have greater connectivity (i.e., they are hubs). Positive and negative correlations between nodes are represented by the edge color blue and red respectively.

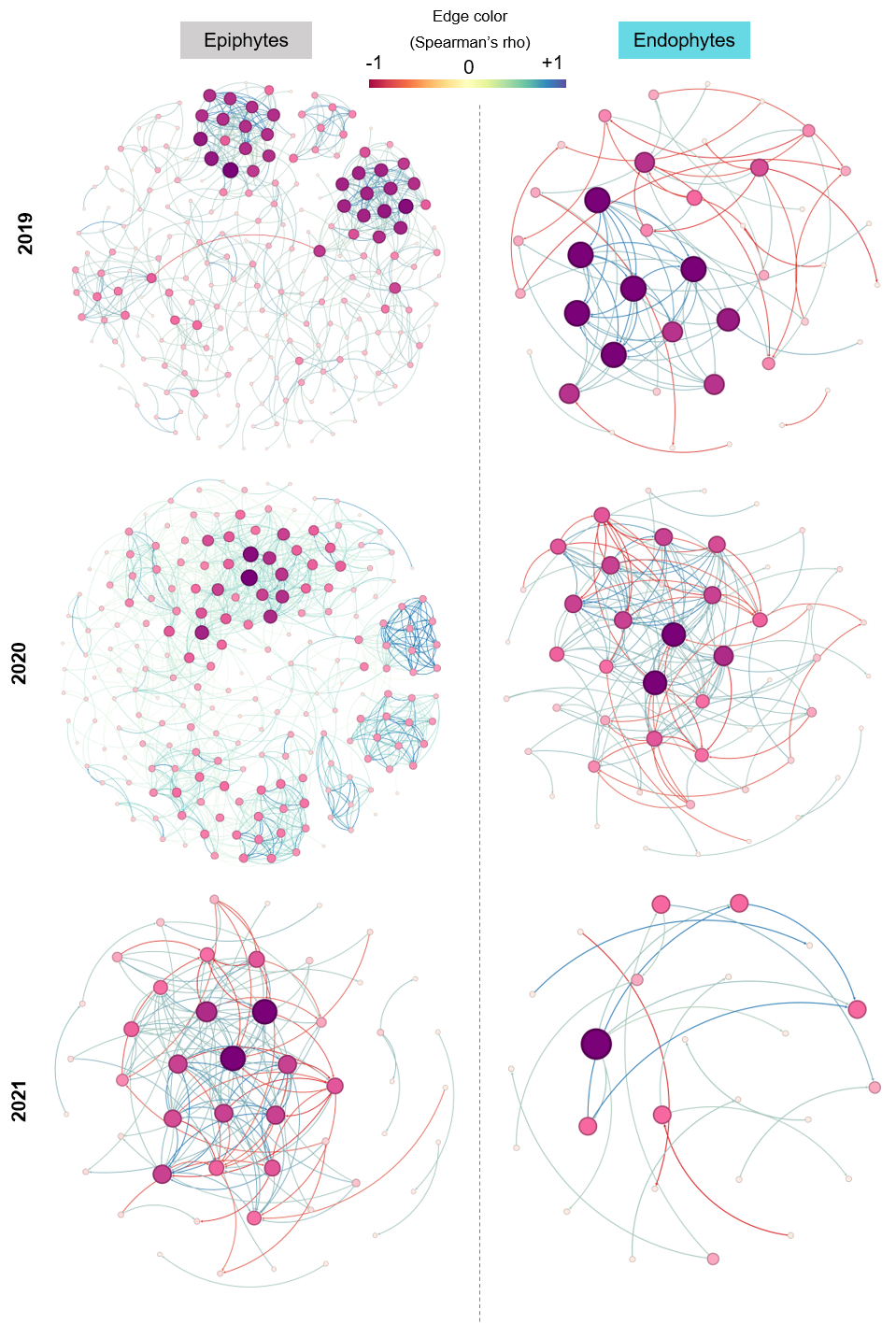

**Fig S9: Additional analysis of co-occurrence networks.** (A). The mean proportion of rare taxa (with <1% of total reads) across years and landraces. Error bars represent the standard error from 4-10 biological replicates/landraces/year/community type. (B). Co-occurrence networks of dominant taxa, excluding 2019 samples from the landrace Chakhao.

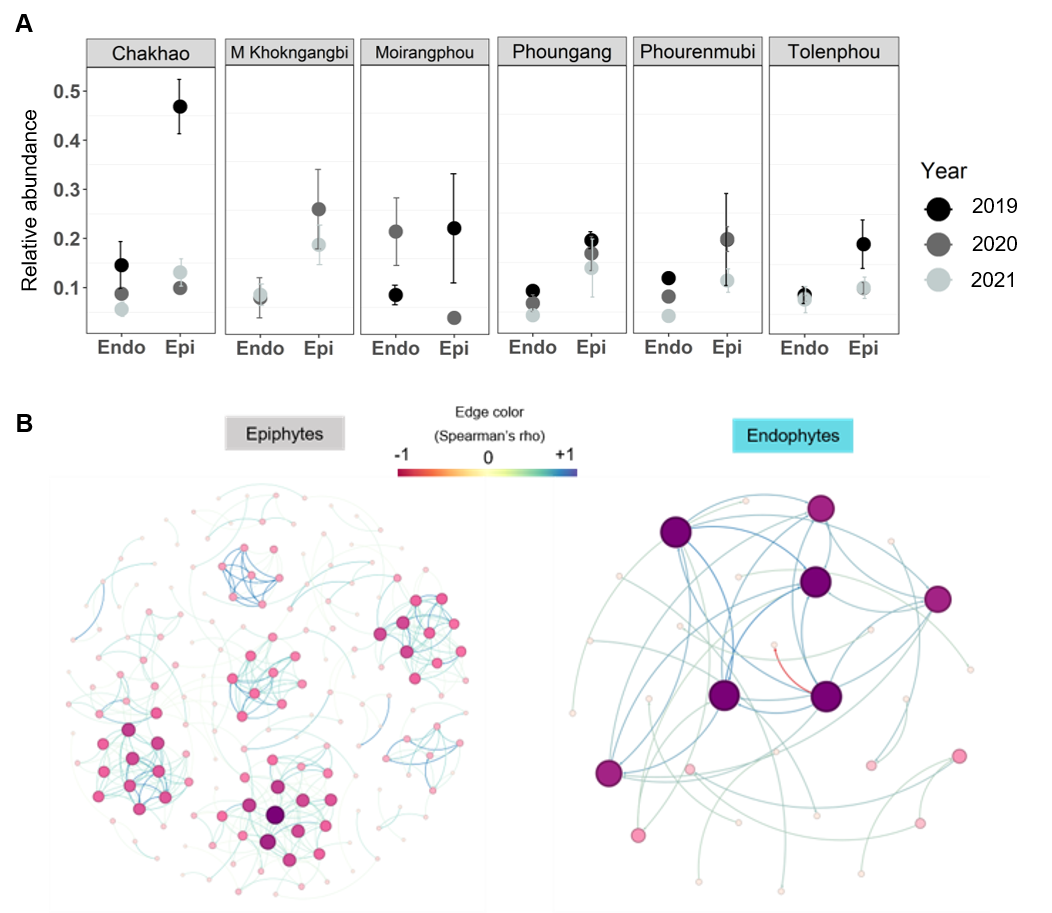

**Fig S10: Correlations between the relative abundance of *Methylobacterium* and other bacterial genera.** Bubble plots showing pairwise correlations between the relative abundance of *Methylobacterium* with other bacterial taxa across years and landraces in (A) epiphytic and (B) endophytic communities. Larger bubbles indicate larger values of Spearman’s rho correlation coefficient. Dotted horizontal grey lines indicate taxa with a consistently strong correlation across all three years.

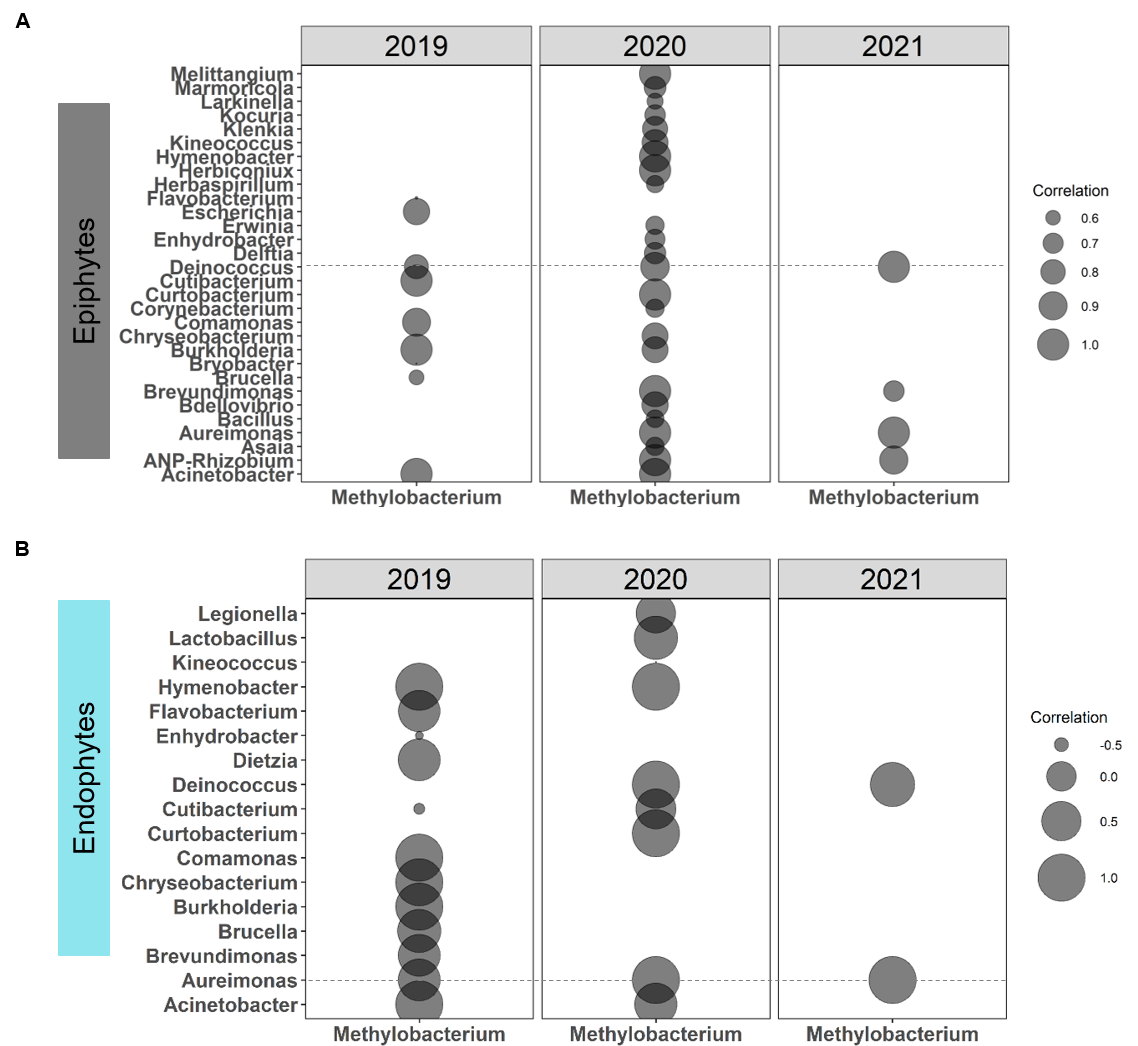

**Fig S11: Epiphytic and endophytic co-occurrence networks of focal landraces.** Epiphytic and endophytic networks of (A) Chakhao, (B) Phouren-mubi, (C) Phoungang, and (D) Tolen-phou. Nodes represent each bacterial taxon (identified to genus level), node size and color indicate the degree of connection between the nodes. Positive and negative relationships between the nodes are represented by the edge color blue and red respectively.

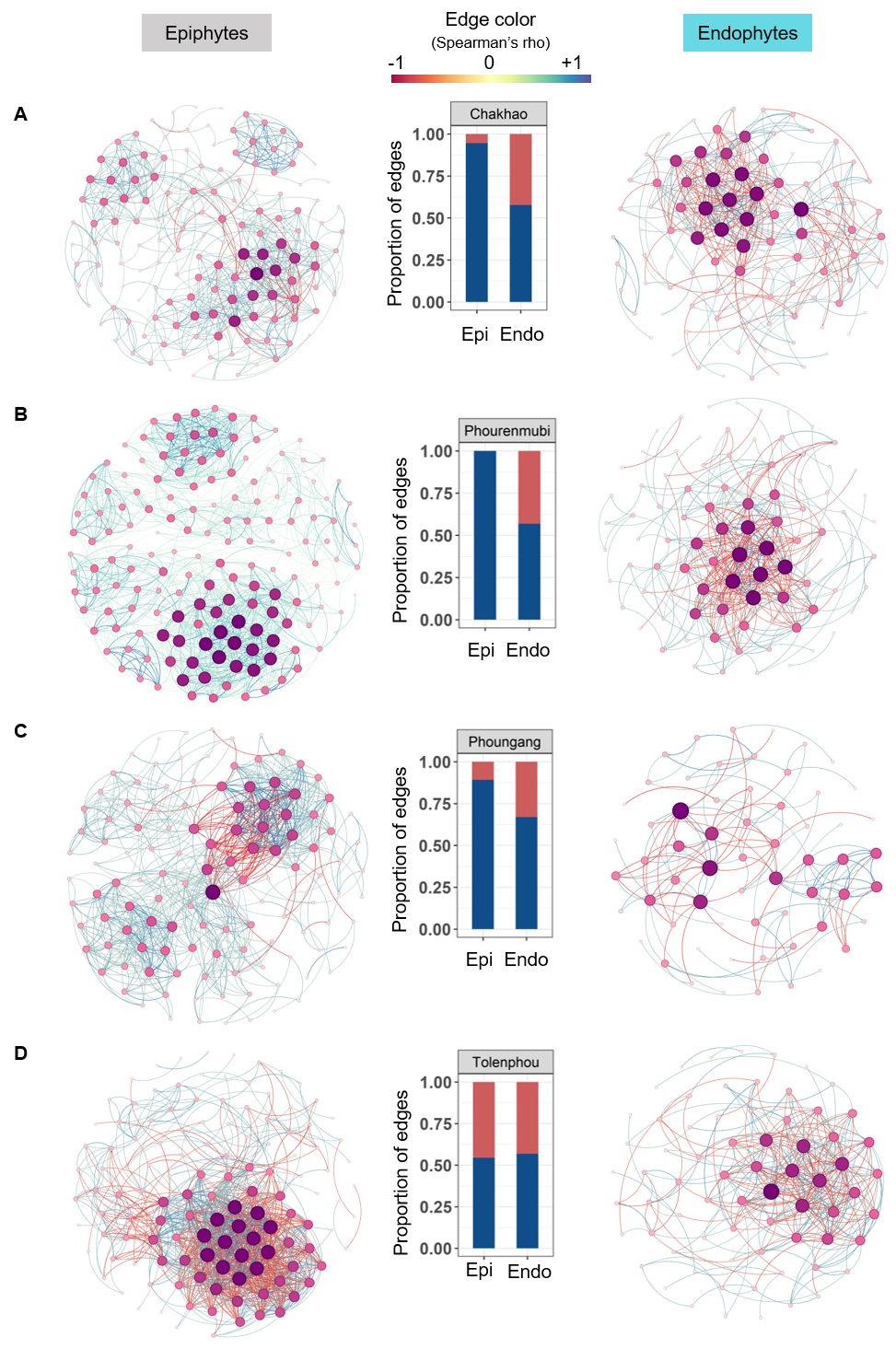

**Fig S12: Richness of phyllosphere microbiomes across growth stages of plants of the landrace Chakhao.** Boxplots show the richness of epiphyte vs. endophyte communities at different growth stages. 10 replicate plants were sampled at each growth stage. Asterisks represent significant differences across communities (paired t-test, P<0.05).

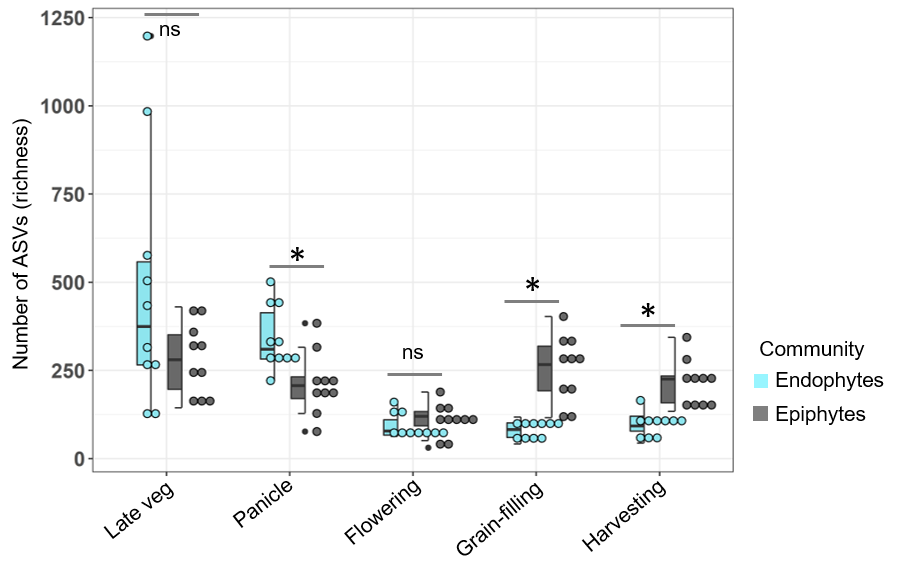

**Fig S13: Differentiation in phyllosphere communities across growth stages of plants of the landrace Chakhao.** (A) Linear discriminant (LD) plots showing the clustering of epiphytic and endophytic bacterial communities across growth stages. (B) Change in the mean proportion of dominant bacterial taxa in both communities, across plant growth stages. Error bars represent the standard error from ten biological replicates/community type/growth stage.

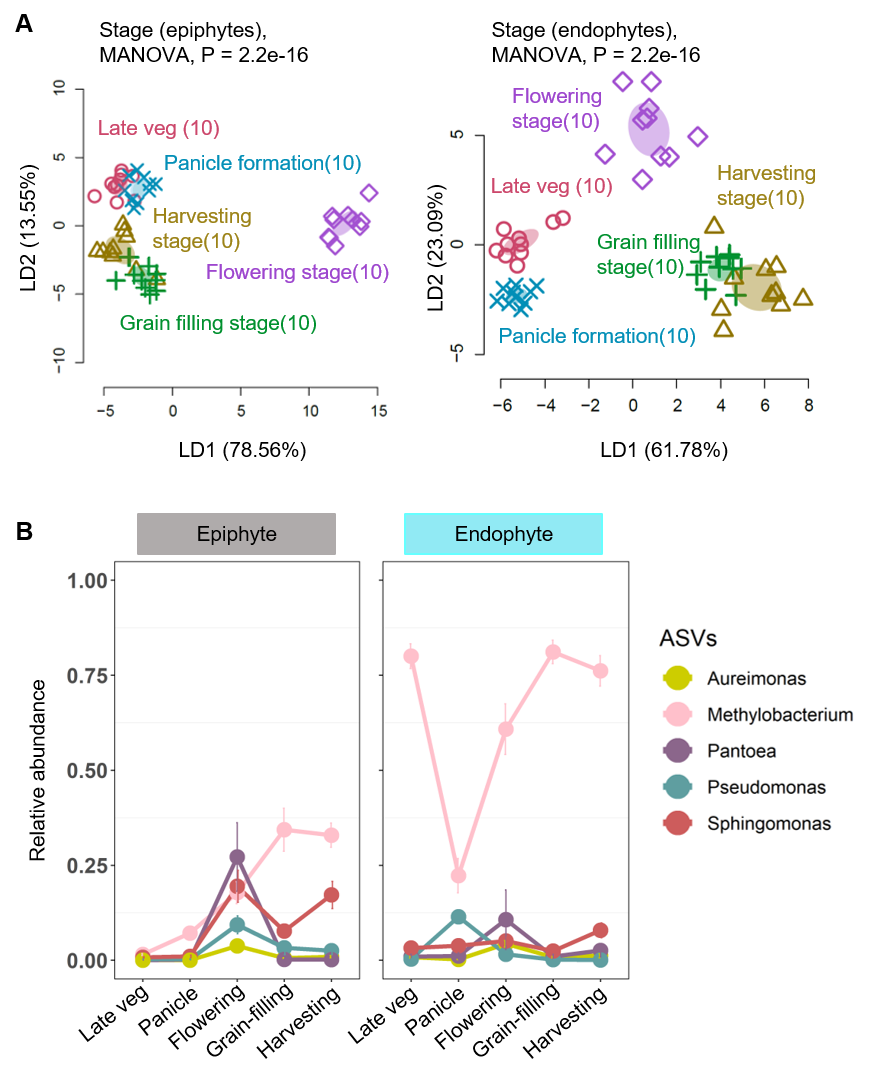

**Fig S14**: **Total bacterial load does not correlate with diversity or richness**. Scatterplots show the relationship between the normalized total bacterial load and Shannon diversity or richness across plant growth stages. Each point represents 5 biological replicate plants/growth stage/community type.

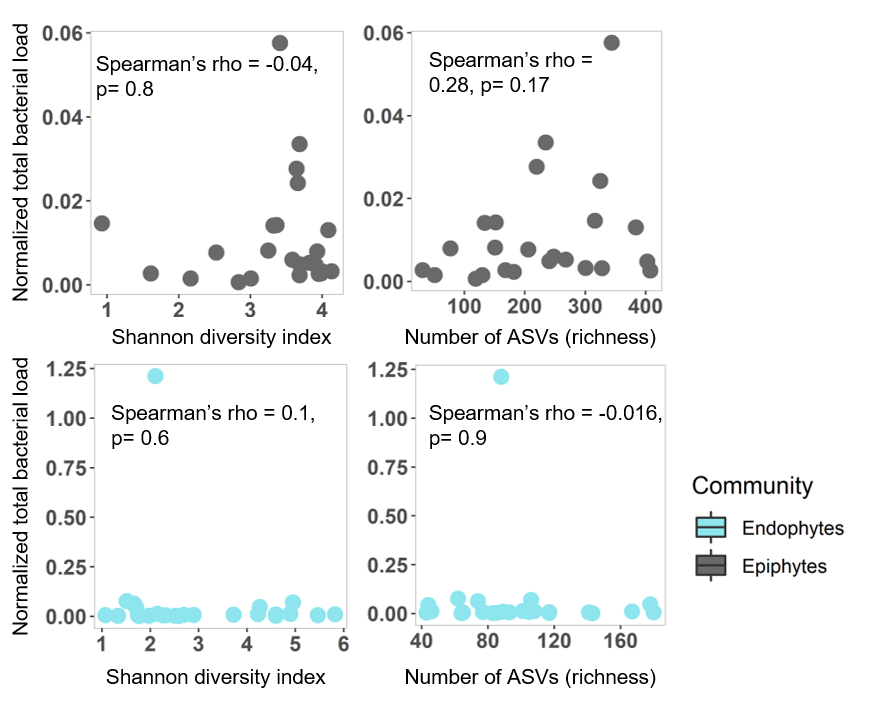
